# Synthetic mycolates derivatives as molecular tools to decipher protein mycoloylation, a unique post-translational modification in bacteria

**DOI:** 10.1101/2024.03.28.587066

**Authors:** Emilie Lesur, Yijie Zhang, Nathalie Dautin, Christiane Dietrich, Ines Li de la Sierra-Gallay, Luis Augusto, Paulin Rollando, Noureddine Lazar, Dominique Urban, Gilles Doisneau, Florence Constantinesco-Becker, Herman Van Tilbeurgh, Dominique Guianvarc’h, Yann Bourdreux, Nicolas Bayan

**Affiliations:** Université Paris-Saclay, CNRS, Institut de Chimie Moléculaire et des Matériaux d’Orsay (ICMMO), UMR 8182, F-91405 Orsay, France; Université Paris-Saclay, CEA, CNRS, Institute for Integrative Biology of the Cell (I2BC), 91198, Gif-sur-Yvette, France

**Author notes:** Université Paris Cité, CNRS, Biochimie des Protéines Membranaires, F-75005 Paris, France. Both authors equally contributed to this work (Emilie Lesur, Yijie Zhang).

## Abstract

Protein mycoloylation is a newly characterized post-translational modification (PTM) specifically found in *Corynebacteriales*, an order of bacteria that includes numerous human pathogens. Their envelope is composed of a unique outer membrane, the so-called mycomembrane made of very-long chain fatty acids, named mycolic acids. Recently, some mycomembrane proteins including PorA have been unambiguously shown to be covalently modified with mycolic acids in the model organism *Corynebacterium glutamicum* by a mechanism that relies on the mycoloyltransferase MytC. This PTM represents the first example of protein *O*-acylation in prokaryotes and the first example of protein modification by mycolic acid. Through the design and synthesis of trehalose monomycolate (TMM) analogs, we prove that *i*) MytC is the mycoloyltransferase directly involved in this PTM, *ii*) TMM, but not TDM, is a suitable mycolate donor for PorA mycoloylation, *iii*) MytC is able to discriminate between an acyl and a mycoloyl chain *in vitro* unlike other trehalose mycoloyltransferases. We also solved the structure of MytC acyl-enzyme obtained with a soluble short TMM analogs which constitutes the first mycoloyltransferase structure with a covalently linked to an authentic mycolic acid moiety. These data highlight the great conformational flexibility of the active site of MytC during the reaction cycle and pave the way for a better understanding of the catalytic mechanism of all members of the mycoloyltransferase family including the essential Antigen85 enzymes in *Mycobacteria*.

## Introduction

*Corynebacteriales* are an order of diderm actinomycetes. Their envelope is constituted of an inner membrane and a thick peptidoglycan-arabinogalactan (PG-AG) polymer on which a specific outer membrane is covalently anchored. All members of *Corynebacteriales* produce long α–ramified and β-hydroxylated fatty acids called mycolic acids (*1*). These unique molecules are produced in the cytoplasm by condensation of two fatty acid molecules that are immediately esterified on trehalose by a polyketide synthase (Pks13) (*2*, *3*). The resulting trehalose monomycolate (TMM) is then translocated across the inner membrane by dedicated Resistance-Nodulation-Division (*RND*) family transporters (*4*) and serves as a donor of mycolate chain for various envelope acceptors such as TMM itself to form trehalose dimycolate (TDM) or, alternatively, for some arabinosyl terminal ends of arabinogalactan (AG). The resulting mycoloylated external surface of the AG polymer will constitutes the outer membrane inner leaflet while TDM together with some other specific lipids will form the outer leaflet of this peculiar membrane, also called the mycomembrane (*5*). The reactions of mycolate transfer from TMM to various acceptors are known to be catalyzed by mycoloyltransferases (Myts) (*6*), a family of enzymes specifically present in all members of *Corynebacteriales*. Several paralogs of Myts coexist in a given species but their exact function/specificity *in vivo* is still to be determined. While six mycoloyltransferases are described in *C. glutamicum*, MytA and MytB are the most expressed in cell laboratory conditions (*7*, *8*). *In vivo*, they have partially redundant functions since they are able to partially replace each other to some extent in TMM and arabinose mycoloylation.

The mycomembrane is unique to *Corynebacteriales* and reminiscent of the lipopolysaccharide (LPS) outer membrane of Gram-negative bacteria. It represents a hydrophobic barrier protecting the cell from noxious hydrophilic compounds and its permeability to small solutes is strictly determined by porins, which have been described in several genera of *Corynebacteriales*. In fast growing mycobacteria, MspA is the main porin so far described and its structure is clearly related to classical β-barrels (*9*) of LPS containing outer membranes. In slow growing bacteria, the proteins from the PE/PPE family seem to fulfill this function, although their mycomembrane-embedded structure is still unknown (*10*). Finally, in *Corynebacteria*, small channel forming proteins have been described but their structure and their function *in vivo* are still very poorly documented (*11*, *12*). In *C. glutamicum* PorAH and PorBC account for a cationic and an anionic porin respectively with estimated conductances of about 2.5 nS and 0.7 nS. Mutants deleted for PorA/PorH and grown on citrate, which blocks PorBC porin, are affected for growth. PorAH is constituted of two small polypeptidic chains of 45 (PorA) and 57 amino acids (PorH), both devoid of any signal sequence (Twin Arginine Translocation (TAT) or general Secretion (Sec) pathways). PorBC is composed of PorB and PorC homologous polypeptidic chains of about 130 amino acids. Monomeric PorB appears to have a helical structure (*13*) which could form a pentamer in the membrane, but there is no experimental structure available so far for this putative complex. Interestingly, PorA, PorB, PorC and PorH have been proved to be post-translationally modified by one or several mycolate chains on dedicated serines (*14*, *15*). Although important for membrane association, the function of this modification remains elusive for these transmembrane proteins. The pore forming activity of PorA/PorH *in vitro* is abolished if PorA is not mycoloylated while membrane insertion of the protein does not seem to be affected (*16*). On the contrary, PorH and PorBC association to the membrane is largely affected in the absence of mycoloylation (*17*, *18*). Porin mycoloylation is abolished in a mutant strain deleted for MytC, one of the six mycoloyltransferases described in *C. glutamicum* strongly suggesting that this protein is somehow involved in the process, either directly or indirectly. In this study, we performed an *in vitro* study of protein mycoloylation using synthetic TMM and appropriately designed TMM analogs. In particular, we definitely showed that MytC was sufficient on its own to carry out PorA mycoloylation and we characterized the crucial structural elements of the mycolate donor, necessary for the transesterification reaction catalyzed by MytC.

## Results

### MytC is able to mycoloylate PorA *in vitro*

*In vivo*, deletion of MytC in *C. glutamicum* is associated with the absence of PorA, PorH and ProtX *O*-mycoloylation indicating that MytC is clearly involved in this uncommon post-translational modification (*19*). In order to confirm that MytC alone is directly able to mycoloylate a protein, we decided to set up an *in vitro* test to evaluate the ability of purified MytC (*19*) to transfer a mycolate moiety from a lipid donor to a protein acceptor. In a first approach we decided to test a mycomembrane lipid extract (highly enriched in TMM and TDM and hereinafter referred to as TDM/TMM mix (see supplementary Figure 1b), as a potential mycolate donor and purified non-mycoloylated PorA as the final acceptor of the reaction (Figure 1a). The mycoloylation of PorA was followed by SDS PAGE since mycoloylated and non-mycoloylated PorA proteins are clearly resolved on a 16% Tris-Tricine SDS-PAGE (*16*). As shown in Figure 1b, when non-mycoloylated PorA (purified from a Δ*mytC* strain) is incubated in the presence of TDM/TMM mix and MytC, an additional band migrating similarly to PorA purified from a wild-type strain (mycoloylated PorA) is detected. This band is absent if MytC is replaced in the reaction mixture by a variant that carries a mutated catalytic serine (S189V) excluding that the observed activity is due to a trace of contaminating protein. As expected, MytA, which is known to be involved in TMM and arabinogalactan mycoloylation but the absence of which does not suppress PorA mycoloylation *in vivo*, is not able to mycoloylate PorA in similar conditions.

**Figure 1.**
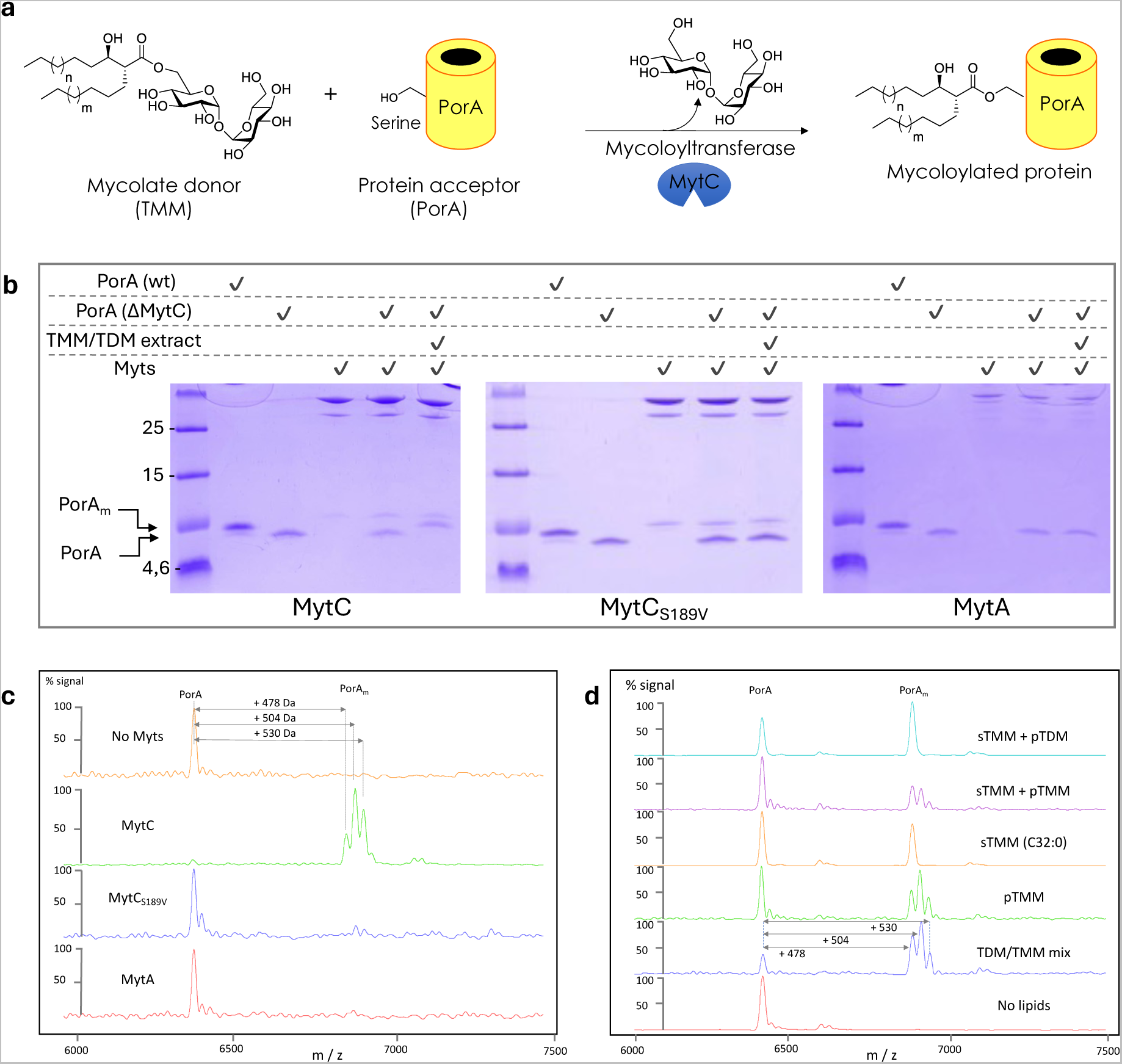
Mycoloylation of PorA *in vitro* by purified MytC. (a) Hypothetical reaction proposed for MytC. (b-d) Analysis of PorA mycoloylation *in vitro* (b) by Tricine-SDS/PAGE and visualized by Coomassie blue coloration; (c-d) by MALDI-TOF MS spectra of the PorA substrate (*m/z* 6410), (c) leading to the formation of the expected major mycoloylated products (*m/z* 6888 (C32:0), 6914 (C34:1), 6940 (C36:2)) in the presence of MytC but not with MytCS189V or MytA, (d) leading to the formation of the expected mycoloylated products in the presence of TMM, purified from bacteria (pTMM) or synthetic (sTMM) but not TDM.

In order to definitively confirm that mycoloylated PorA is generated during the reaction, all samples were analyzed in parallel by mass spectrometry (Figure 1c). In the presence of MytC, but not with the null variant of MytC nor with MytA, the peak corresponding to native non-mycoloylated PorA (6410 Da) shifted to a series of three main peaks with a mass difference of 478, 504 and 530 Da compared to the mass of unmodified PorA. These peaks perfectly match the natural occurrence of the three major mycolic acid species that have been described in *C. glutamicum* (C32:0, C34:1 and C36:2) (*2*, *14*). Altogether these results unambiguously demonstrate that purified recombinant MytC is able to mycoloylate PorA when a mixture of mycomembrane lipids, i.e. TDM/TMM mix, is provided as mycolate donors.

### TMM, but not TDM, is a suitable mycolate donor for PorA

Only sparse information is available concerning the mechanism involved in protein mycoloylation *in vivo*, and more particularly little is known about the nature of the donor substrate. In a previous study, we showed that TMM accumulated in a Δ*mytC* strain, suggesting that this molecule could be the natural mycolate donor (*15*). However, TDM, the other major trehalose mycolate ester found in the cell envelope may also potentially constitute a suitable donor for MytC. Alternatively, minor glycolipids present in the mycomembrane could also be used by MytC. To address this question, we first decided to test either purified or synthetic TMM molecules as substrates in the reaction. Purified TMM (pTMM) was prepared from our bacterial mycomembrane extract as described in the Materials and Methods section and synthetic TMM (sTMM) was obtained as previously described (*20*). Both preparations (see Supplementary Figure 1) of TMM are easily resuspended in aqueous solution and, as shown in Figure 1d, are readily used by MytC to mycoloylate PorA. Indeed, when either a mycomembrane lipid extract (TDM/TMM mix) or purified TMM are used in the reaction, the three characteristic main peaks corresponding to mycoloylated PorA are detected in both conditions. Instead, when synthetic C32:0 TMM is used, only a single peak corresponding to PorA modified by a C32:0 mycolate chain (Δm = 478 Da) is detected. These results clearly show that TMM is a *bonafide* donor of mycolate for protein mycoloylation *in vitro*. Because natural or synthetic TDM are mostly insoluble and very difficult to resuspend correctly in aqueous solution (*21*), these substrates cannot be used for an *in vitro* MytC assay. We decided to resuspend the TDM in aqueous solution by introducing synthetic TMM C32:0 in our TDM preparations (ratio 1:1). In these conditions, we noticed that the TMM-C32:0/TDM lipid film is nicely rehydrated and does not form any visible large insoluble material. Interestingly, when this mixture is further used as a mycolate donor for PorA mycoloylation by MytC in our *in vitro* test, we clearly only observe the appearance of a single modified PorA species corresponding to the transfer of the C32:0 chain (Δm = 478 Da) but not of the expected C34:1 or C36:2 chains that would have appeared if purified TDM had been used as a donor. As a control, using a mixture of synthetic C32:0 TMM and purified TMM, we were clearly able to detect the characteristic series of three peaks corresponding to PorA species modified by all major mycolate chains (C32:0, C34:1 and C36:2) of “natural” TMM. Altogether, these results strongly suggest that TDM is probably not an efficient donor of mycolate for PorA mycoloylation.

### Chemical synthesis of TMM analogs

To further define the specificity of MytC towards its mycolate acceptors, we decided to chemically synthesize several TMM analogs with modifications introduced on the mycolate moiety (Figure 2a). They were synthesized either with shorter branched acyl chains (compounds **1** (TMM-C23:0) and **2** (TMM-C13:0)), or lacking the hydroxyl group (compound **3**), lacking the α-alkyl chain (compounds **4** and **5**), lacking the H bond donor capacity of the hydroxyl group (compound **5**), or only composed of a simple unsubstituted acyl chain (compound **6**, also named trehalose monopalmitate **(TMP)**). To date, most TMM-based reported tools contain simplified fatty acyl chain thus making their syntheses fast and efficient (*22*). Recently, we reported the first synthesis of a bioorthogonal TMM-based probes with the natural pattern of mycolic acid which proved to be highly efficient in labeling *C. glutamicum* (*23*). Based on these results, we anticipated that TMM derivatives should help to decipher the enzymatic mechanism and specificity of MytC. Such compounds represent an important synthetic challenge to include the native pattern of mycolic acids featuring an OH group in β-position with an (*R*) stereochemistry and a branched α-lipid chain with an (*R*) stereochemistry and an *anti*-relationship at C_2_-C_3_ between the two β and α-substituents. Our TMM library was synthesized through two main steps: i) preparation of the fatty acids parts based on Noyori enantioselective reduction of β-ketoester (*24*) and diastereoselective alkylation of the resulting β-hydroxyester using Fráter-Seebach alkylation (*25*, *26*) and ii) esterification on a selectively protected trehalose derivative (Figure 2a). Figure 2c shows a brief overview of the TMM derivatives synthesis. For non-ramified and non-hydroxylated TMM analog **6**, commercially available fatty palmitic acid **7b** was used and esterified to trehalose **15**. For TMM analog **3**, carboxylic acid **8** was obtained by α-alkylation of **7b** with 1-iodohexadecane. Derivatives **9a-c** were obtained after a two-atom homologation of their corresponding carboxylic acids **7a-c**. Derivatives **9a-c** were subjected to enantioselective Noyori reduction to afford the β-hydroxyesters **10a-c**. Derivative **10b** was then converted to carboxylic acid **11** which was then esterified with selectively protected trehalose **15** to give TMM analog **4**. **10b** was also treated with proton sponge® and trimethyloxonium tetrafluoroborate followed by aqueous sodium hydroxide to furnish carboxylic acid **12** which was coupled to trehalose **15** to give analog **5**. Compounds **10a** and **10c** were finally treated with LDA (lithium diisopropylamide) and iodoalkane (*i.e.* 1-iodopentane or 1-iododecane) to furnish derivatives **13a** and **13b** which were then converted to carboxylic acids **14a** and 1**4b**, and further esterified with trehalose **15** to give TMM analogs **1** and **2.**

**Figure 2.**
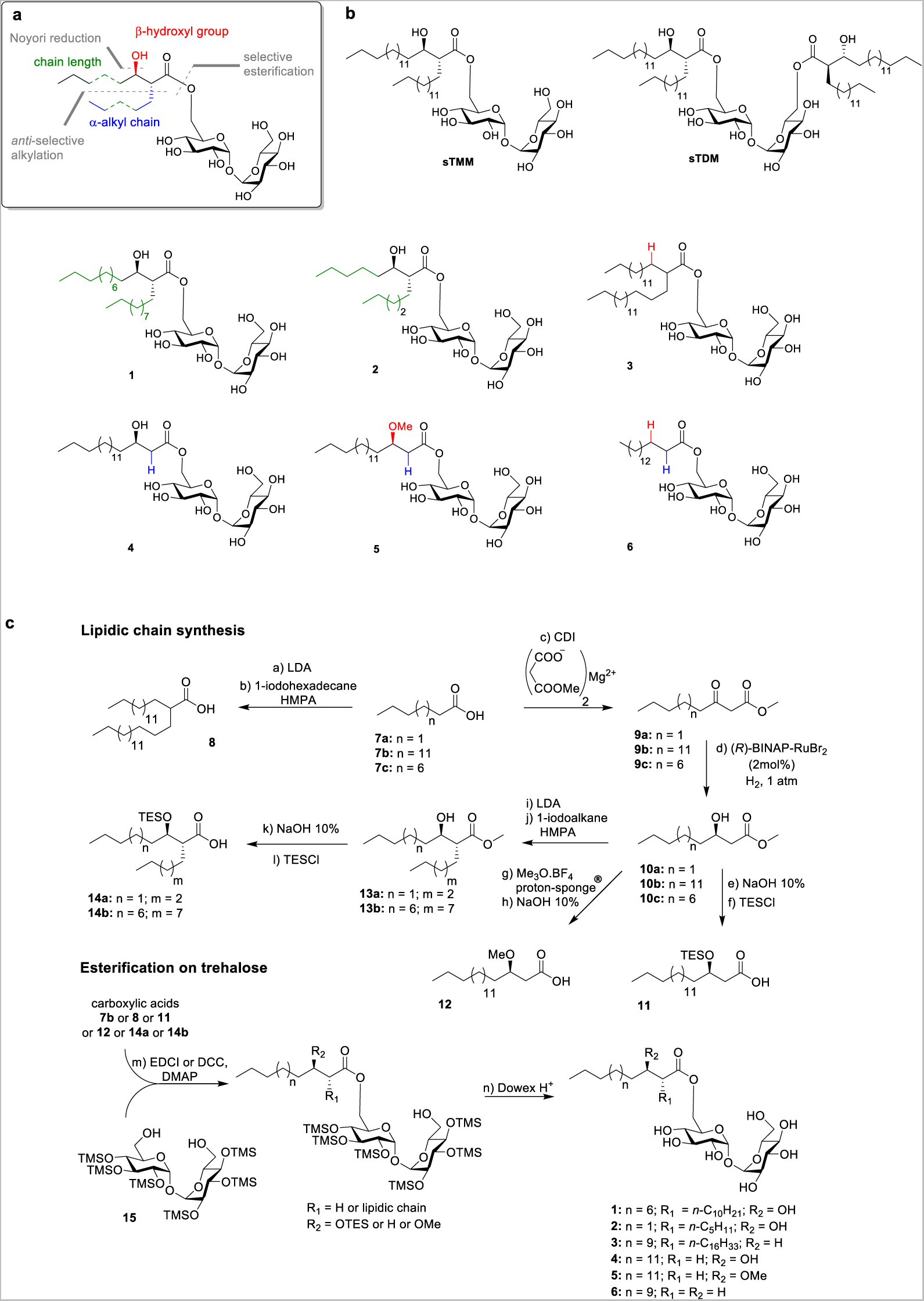
Design and synthesis of a TMM analogs library. (a) Synthetic strategy to obtain the TMM analogs with a mycolate pattern featuring the stereochemistry of the natural compounds. (b) Synthetic “natural” TMM and TDM (C32:0) and new TMM analogs. The colors highlight the modified part of the mycolate pattern according to the defined code in (a); (c) brief overview of the TMM derivatives synthesis.

### The β-hydroxylation of the mycolic acid chain of TMM is an essential determinant of MytC specificity

To evaluate the substrate specificity of MytC, we first assessed the ability of these TMM analogs to serve as mycolate donor. For all of them we assayed esterification (mycoloylation or acylation) of PorA in the presence of MytC in conditions strictly identical to those used in the assay described above. Indeed, we included for all experiments a control of PorA mycoloylation with the TDM/TMM mix in parallel. Because some analogs are not very soluble (see above for TDM), we systematically tested the different compounds alone or in combination with the TDM/TMM mix (ratio 1:1 between analog and TMM of the mix) in order to provide more homogeneous and comparable preparations of mycolate donor. The samples were only analyzed by mass spectrometry because the modifications of PorA by some of the very short synthetic chains would not have been resolved on SDS-PAGE.

When PorA is incubated in the presence of analog **1** (synthetic mycolate C23:0) alone or in combination with the TDM/TMM mix, a major peak at 6758 Da is obtained, corresponding to PorA modified by a C23:0 mycoloyl chain (Δm = 352 Da) (Figure 3a, Table 1). The much shorter trehalose mycolate analog **2** (synthetic mycolate C13:0), was able to modify PorA by a C13:0 mycolate chain (peak at 6610 Da, Δm =212 Da) only in combination with the TDM/TMM mix (Figure 3a, Table 1). This indicates that analog **2** may form micelles or other specific structures that are less accessible to MytC than those formed when combined with TDM and TMM liposomes.

**Figure 3.**
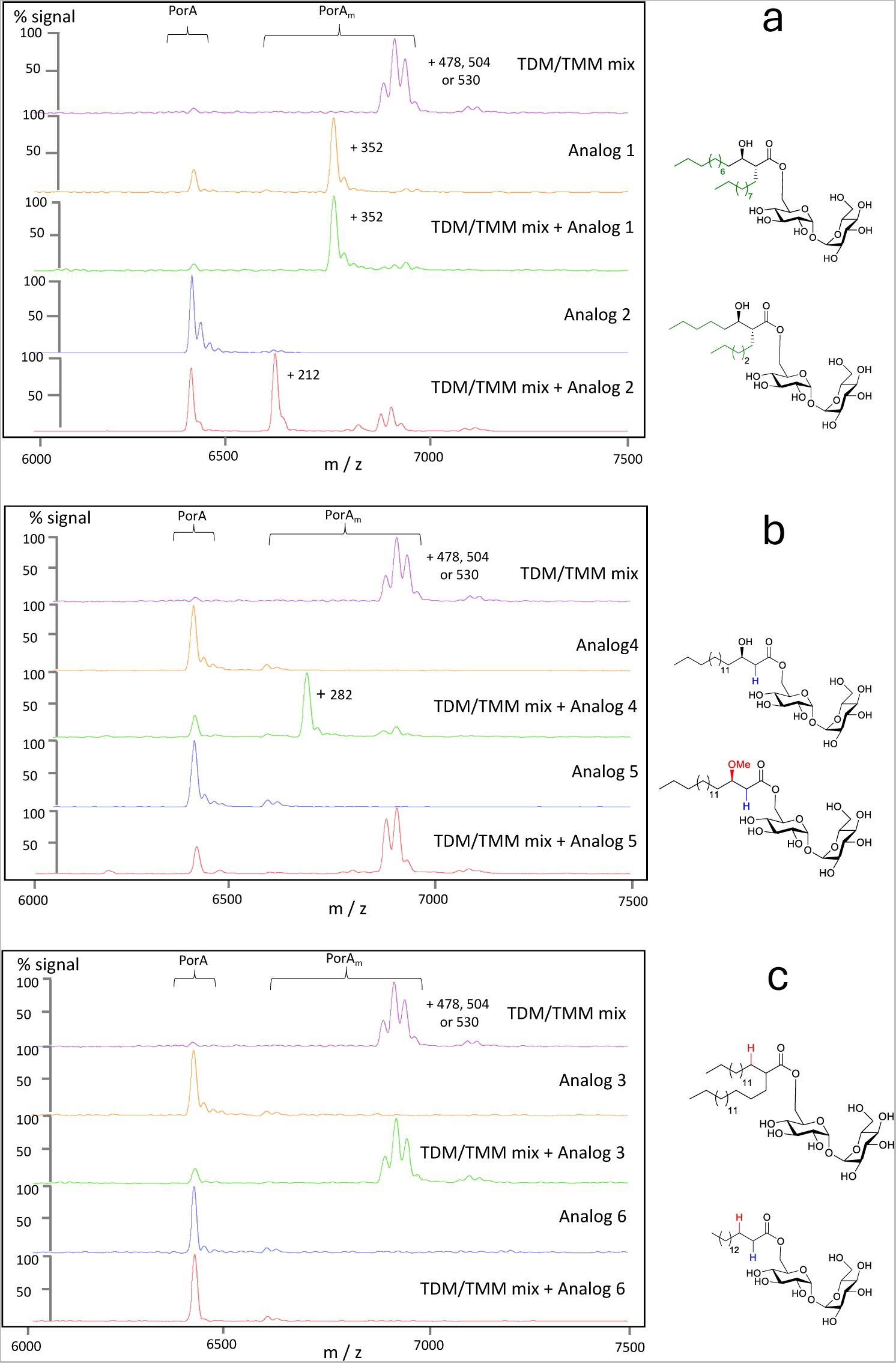
Mycoloylation of PorA by MytC using TMM analogs. Analysis of PorA mycoloylation in the presence of MytC and TMM analogs by MALDI-TOF MS.

**Table 1:** Mass of TMM and analogs and expected Δm after fatty acid transfer onto a mycolate acceptor.

|  | TMM or<br>analogs<br>(g/mol) | TMM or<br>analogs<br>(M+Na) | Expected increase in<br>mass after acyl or<br>mycoloyl chain transfer<br>on acceptors |
| --- | --- | --- | --- |
| **C32:0** | 820<br>(820.59) | 843 | + 478.47 |
| **C34:1** | 846<br>(846.61) | 869 | + 504.49 |
| **C36:2** | 872<br>(872.62) | 895 | + 530.51 |
| **Analog 1** | 694<br>(694.45) | 717 | + 352.33 |
| **Analog 2**<br>**(TMM-C13 :0)** | 554<br>(554.29) | 577 | + 212.17 |
| **Analog 3** | 804<br>(804.60) | 827 | + 462.48 |
| **Analog 4** | 624<br>(624.37) | 647 | + 282.27 |
| **Analog 5** | 638<br>(638.39) | 661 | + 296.27 |
| **Analog 6**<br>**(TMP)** | 580<br>(580.35) | 603 | + 238.23 |

Mycolate chains differ from a classical fatty acyl chain by the presence of an alkyl side chain and a hydroxyl group at the α and β positions respectively. To get insight in MytC specificity between acyl and mycoloyl chains, we assayed several analogs of TMM lacking one or both mycolic acid determinants using the same experimental approach as above. Interestingly, the absence of the α-alkyl side chain is not critical for mycoloylation by MytC since analog **4**, which main chain is similar to TMM but lacks the α-branch, is perfectly used as substrate to acylate PorA when provided in combination with TDM/TMM mix (Figure 3b). On the contrary, β-hydroxylation of the mycoloyl chain is clearly essential since compound **3**, an analog of TMM lacking the β-hydroxyl group, is not used as a substrate by MytC in all conditions tested (Figure 3c). Interestingly, the β-hydroxylation of the mycoloyl chain is not only required but also found sufficient to allow the transfer of a single acyl chain on PorA since a β-hydroxylated acyl trehalose is well recognized by MytC (Figure 3b, analog **4**) while a simple acyl trehalose is not (Figure 3c, analog **6**). The β-hydroxyl group of the substrate is probably crucial for hydrogen bonding (either intramolecular or with a critical residue of MytC) since compound **5**, a methoxy version of analog **4** is completely unable to transfer its acyl chain either alone or in combination with the TDM/TMM mix (Figure 3b, analog **5**).

### TMP (Analog 6) is an efficient inhibitor of MytC

The protein mycoloylation reaction proceeds by two distinct steps involving a double transesterification reactions at the catalytic site. In the first step, the mycolate of TMM is transferred on the catalytic serine (S189), forming a covalent mycoloyl-enzyme intermediate, followed by a second transesterification onto the serine of the protein acceptor (Figure 1a). To identify the structural determinants for each step, we assessed the inhibitory capacity of some TMM analogs. Analogs **3** and **6** (TMP) are not used by MytC as substrates since no modification of PorA is detected in their presence. We further tested whether these compounds are inhibitors of MytC. When **3** is used in combination with the TDM/TMM mix, we still observe a modification of PorA by the different mycolate chains of the TMM indicating that it is not recognized by the enzyme. This is not the case for TMP which therefore seems to inhibit the reaction (Figure 3c). To confirm this hypothesis, we incubated PorA and MytC with increasing amounts of TMP with a fixed concentration of TDM/TMM mix (2 mM). As expected, the characteristic peaks corresponding to mycoloylated PorA progressively decreased indicating that TMP is efficiently inhibiting MytC activity as mycoloylated PorA is no more detected at 0.5 and 1 mM of TMP (Supplementary Figure 2).

### Both TMM-C13:0 (analog 2) and TMP (analog 6) are efficient substrates for MytA-catalyzed trehalose mycoloylation

*C. glutamicum* mycoloyltransferases have been shown to transfer mycolate chains on various cell wall acceptors such as trehalose and arabinose in order to build the mycomembrane (*6*). *In vivo*, deletion of MytA has dramatic effects on the synthesis of TDM while deletion of MytC is much less deleterious in glycolipid synthesis (*27*). *In vitro*, mycoloyltransferases of Mycobacteria have been shown to use TMM as a donor substrate to mycoloylate another molecule of TMM giving rise to TDM (*28*). This reaction has been further studied by using various alternative substrates to set up fluoro- or colorimetric assays (*29*, *30*) suitable for high throughput screening. Here we tested the activity of MytA (control) and MytC for their ability to synthesize TDM from synthetic TMM *in vitro* (Supplementary Figure 3). Myts proteins were incubated in the presence of synthetic TMM and the putative formation of TDM was detected by Thin Layer Chromatography (TLC) after sulfuric acid staining or by mass spectrometry. After 3 hours of incubation, a small amount of TDM is detected on TLC when the reaction was performed in the presence of MytA but not in the presence of MytC (Figure 4a). This result is also confirmed by mass spectrometry. In the presence of MytA, a peak corresponding to TDM is clearly detected while only a very small one is detected when MytC is added into the reaction (Figure 4b). These results suggest that, unlike MytA, MytC does not use efficiently TMM efficiently as an acceptor molecule in this reaction. Because we showed that MytC displayed a very strict specificity regarding the mycoloyl and the acyl groups that are transferred on PorA, we checked whether MytA has the same requirements in its ability to mycoloylate trehalose. For this, we incubated MytA (or MytC as a control) in the presence of analog **2** or **6**. As shown in Figure 4c and 4d, both compounds **2** and **6** are efficient substrates of MytA which is able to transfer either a mycolate or an acyl chain from analog **2** or **6** respectively to another molecule of analog **2** or **6** (Supplementary Figure 3). For MytC, only very small peaks corresponding to the product of the reaction are detected, similarly to what is observed when authentic TMM (C32:0) is used as a substrate. This result indicates that MytA, unlike MytC, is not able to discriminate between an acyl and a mycoloyl chain *in vitro*.

**Figure 4.**
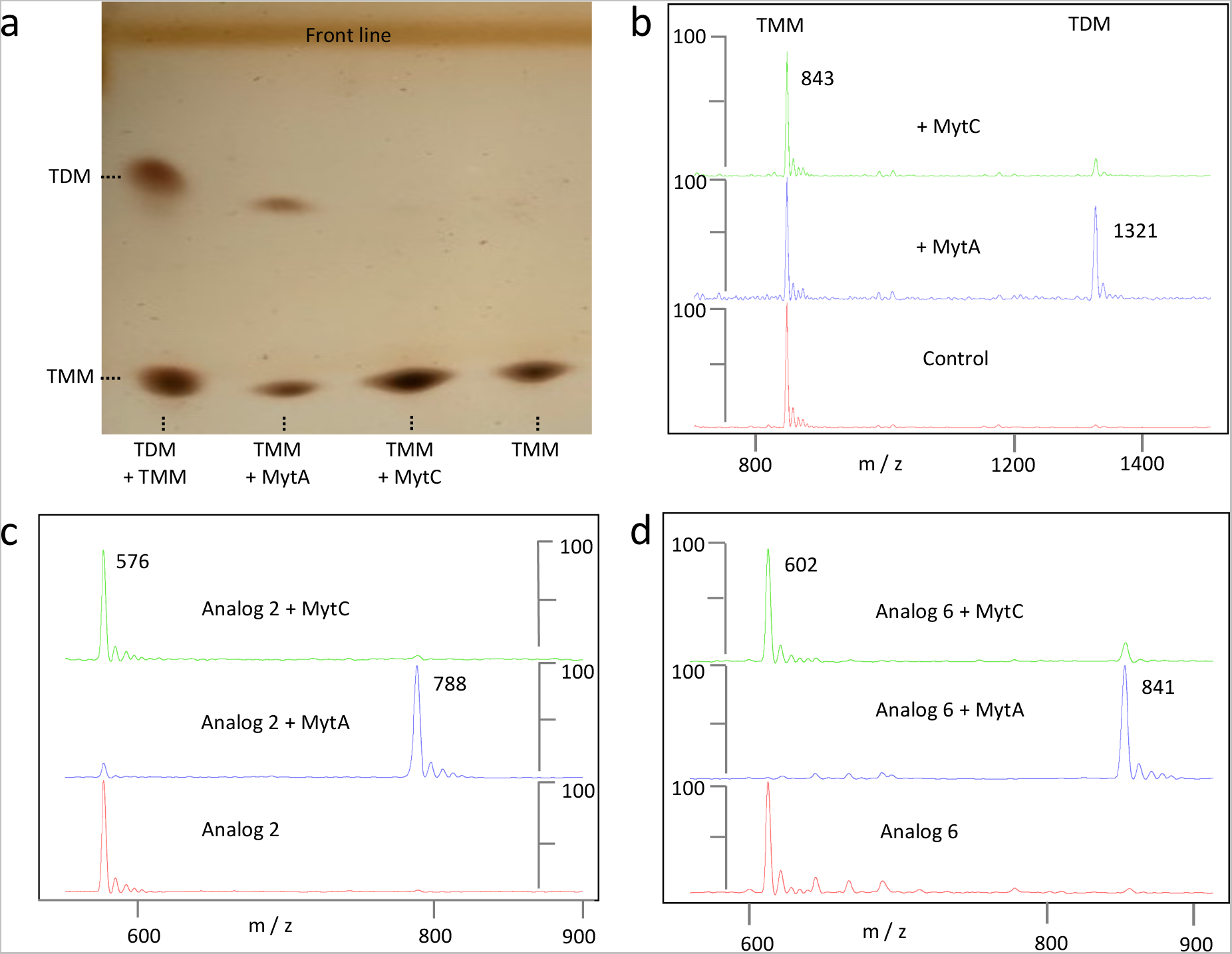
Mycoloylation of TMM and TMM analogs by MytA and MytC. (a and b) Analysis of sTMM mycoloylation in the presence of MytC and MytA by TLC (a) or MALDI-TOF MS (b). (c and d) Analysis of TMM analogs **2** and **6** mycoloylation by MALDI-TOF MS. The scale represents the % of the signal.

### TMM-C13:0 (analog 2) traps MytC in its mycoloyl-enzyme transition covalent intermediate

The resolution of enzyme structures in the presence of mycolic acid-like substrate has significantly increased the understanding of the reaction mechanism of mycobacterial mycoloyltransferases. For instance Ag85C has been crystallized in the presence of diethyl p-nitrophenyl phosphate (*31*), octyl-glucoside (*32*) and more recently with tetrahydroxylipstatin (THL) yielding a covalent enzyme inhibitor complex (*33*). Based on these structures a convincing catalytic mechanism consisting of two successive mycolate transfer reactions could be proposed. Here, to further explore the mechanism and the specificity of protein mycoloylation, we took profit of our TMM analog library to obtain crystals from MytC trapped in its acyl enzyme state. We tested both the TMM-C13:0 (analog **2**), a water-soluble analog of TMM and TMP, the previously characterized inhibitor of the reaction. We obtained diffracting crystals of MytC with both compounds and their structure were solved by molecular replacement. TMP was not detected in the crystal and the structure of MytC determined in these conditions was identical to the one obtained for the apoenzyme. The structure of the crystal of MytC incubated with TMM-C13:0 was solved at 2.69Å (PDB accession code 8QHF) (Figure 5a and 5b). The space group was F23 and the asymmetric unit contained one copy of MytC with a solvent content of 49% (see Supplementary Table 1). The Fo-Fc omit map shows that S189 Oγ atom forms an ester bond with the carbonyl of the analog (Supplementary Figure 4). This structure will be further referred to as the MytC-acyl enzyme (MytC with catalytic S189 esterified). Electron density in the MytC-acyl enzyme was well defined for residues 30 to 363, except for the segment between 283 to 287. The meromycolic chain of the mycolate C13:0 is directed towards the exit of the active site cave, while the α-chain is more deeply buried and establishes hydrophobic interactions with residues of MytC α11 helix (Supplementary Figure 5a). The β-hydroxyl group, a determinant for PorA modification, does not establish polar interactions with MytC. Modeling of full-length alkyl chains of a TMM substrate (C32:0) onto the MytC-acyl structure shows that these could be comfortably accommodated by the MytC active site canyon (Supplementary Figure 5a and 5b). Interestingly, comparison of the active sites of MytC-acyl enzyme and Ag85C-THL (5VNS) (Supplementary Figure 5c) shows that, although the main chain atoms of the catalytic serines (S189 and S124 respectively) superpose exactly, the dihedral χ^1^ angles of the side chains differ by about 60°, and the ester-carbonyl of the two covalent intermediates are in totally opposite directions. While the carbonyl in the tetrahydrolipstatin intermediate points towards a putative oxyanion hole, the equivalent carbonyl of MytC-acyl enzyme does not establish polar interactions with the enzyme. The β-OH groups occupy about the same position in both enzymes despite the opposite configuration at the carbon C-3 (Supplementary Figure 5d).

**Figure 5.**
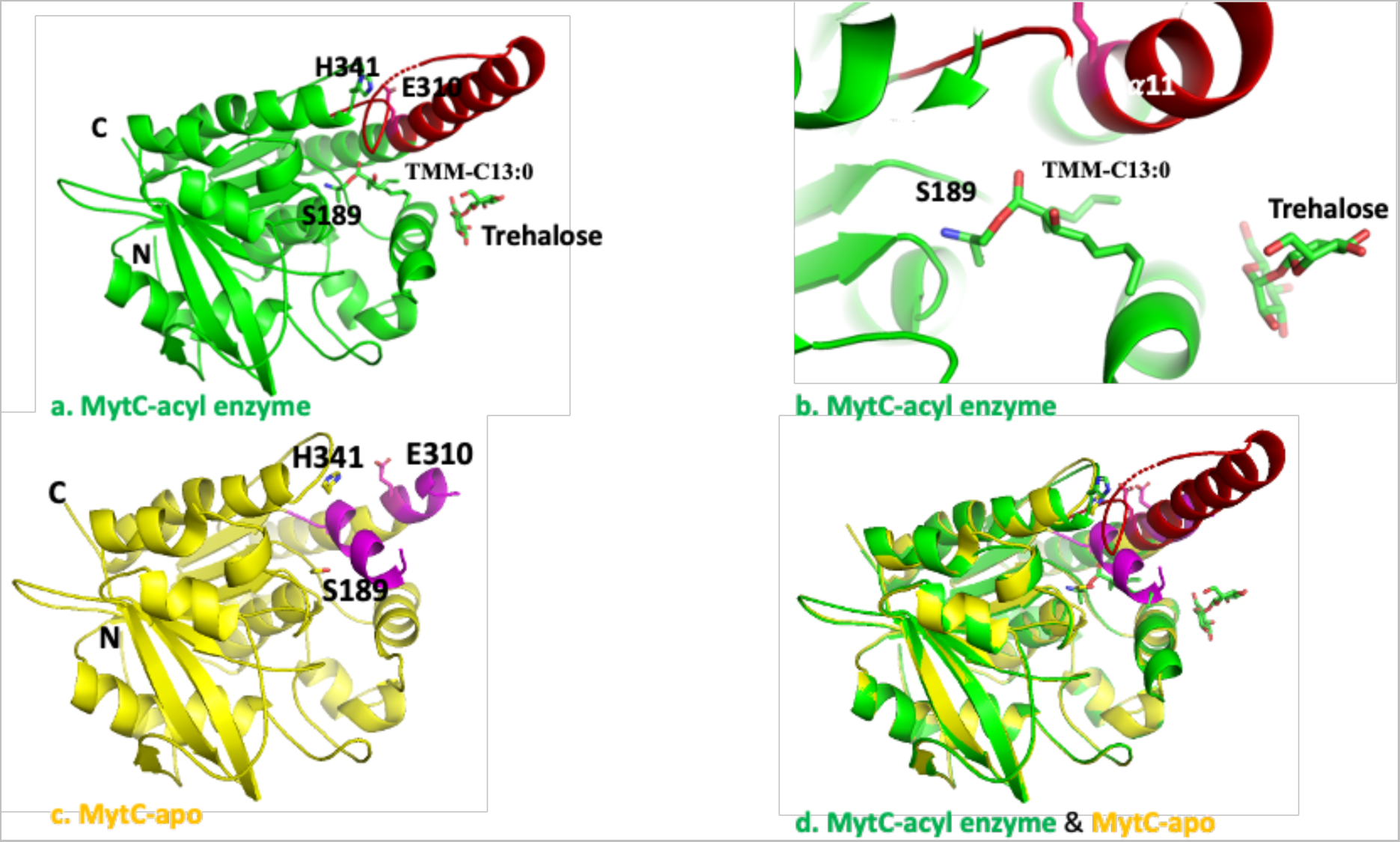
Crystal structure of the MytC-acyl enzyme. (a) Ribbon presentation of of the MytC-acyl enzyme. The *N* and *C* termini are labelled as well as the residues composing the catalytic triad (S189, E319 and H341). Covalently bound TMM-C13:0 and trehalose are in sticks. The region in red was mostly disordered in the crystal structure of apo-MytC. (b) Zoom of the catalytic site bound to TMM-C13:0. (c) Structure of apo-MytC. The helix blocking the active site and part of the α11 helix are in pink. (d) Superposition of the MytC-acyl and apo-MytC crystal structures.

The MytC catalytic triad residues S189, H341 and E310 are clearly in an inactive configuration: the H341 Nδ is at a distance of 15 Å from the S189 Oγ and 14 Å from the E310 Oε1 (Supplementary Figure 6). As shown in Figure 5c and 5d, the segment between residues 271-311 undergoes a major conformational change in the MytC-acyl enzyme compared to the apo enzyme (PDB accession code 4H18, (*15*)). In the apoform, a small helix (η3) contained between residues 270-280, is positioned between S189, H341 and E310, blocking the active site pocket. This helix unfolds in the MytC-acyl enzyme structure to form an irregular loop, becoming part of the active site wall, thereby providing space for the binding of the α-alkyl moiety of the mycolate chain C13:0. Residues 291 to 304, disordered in the apo form, have a well-defined structure in MytC-acyl enzyme, forming an extension of the α11 helix that contacts the C13:0 alkyl chain.

A trehalose moiety was identified on the surface of MytC, near the entrance of the active site groove. One of the trehalose glucose moiety makes polar contacts with the side chains of residues E225 and D232 of helix α8 (Supplementary Figure 7). The biochemical significance of this binding site remains unclear.

To further investigate the flexibility of the active site region, we compared our MytC-acyl enzyme structure with the alphafold (AF)-model of MytC present in the AF database. The pLDDT score of the AF model is very high (>90) for most of the sequence and is around 70 for the region between residues 272 and 308. The AF-model superposes very well onto the MytC-acyl crystal structure (rmsd of 0.29 Å for 266 Cα positions). The AF model also has a fully formed α11 helix which more pronouncedly bends towards the active site pocket as compared to the MytC-acyl enzyme structure. Interestingly, the AF-model proposes a conformation for the loop between residues 336 and 343 that is different from that of the MytC-acyl enzyme structure. This loop swings into the active site pocket and positions His341 between Ser189 and Glu310. Due to the conformational change of this loop and the bending of the α11 helix, the catalytic triad of the AF model is in a catalytically competent configuration. (Supplementary Figure 6a and 6b).

### Modeling of MytC-PorA interaction by Alphafold

For the moment, there is no experimental structural information available concerning the PorA-MytC interaction. We therefore constructed a model of the MytC-PorA complex using AlphaFold-assembly. The model of the PorA protein alone is of good quality, proposing two helical segments. Interestingly, the targeted Ser15 is situated in a very short linker that connects the two helices (Supplementary Figure 9a). In the MytC-PorA complex, PorA forms a helical hook shaped structure that clips between helices α8 and α11 into the active site of MytC (Figure 6, and Supplementry Figure 9b). Interestingly, the mycoloylation target residue of PorA (S15) is located in a loop that connects two helices of PorA. This loop sits deeply into the active site of MytC facing the catalytic serine (Figure 6). When superposing onto the MytC-acyl structure, the Ser15 Oγ is at 7Å from the carbonyl carbon atom of the mycolate C13:0 (Figure 6, inset). As for the alphafold (AF)-model of apo-MytC, the catalytic residues Ser189, His341 and Glu310 were in an active configuration in the AF-model of the MytC-PorA complex (Supplementary Figure 6b). The superposition also suggests that the β-hydroxyl group of TMM-C13:0 could engage a polar interaction with PorA, which might explain why its presence is required for the mycoylation reaction.

**Figure 6.**
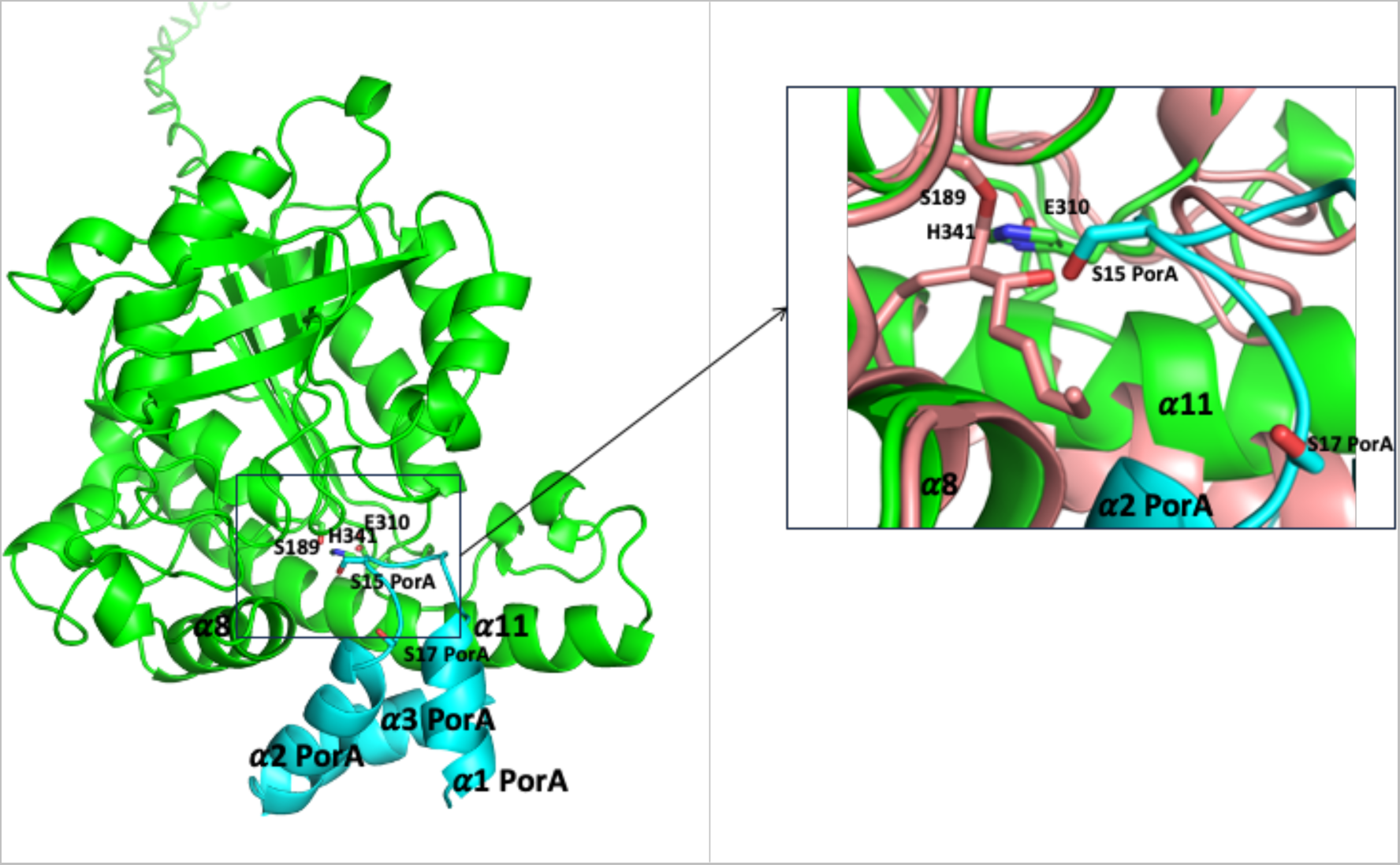
AlphaFold model of the MytC-PorA complex. MytC in green and PorA in light blue. The catalytic triad residues of MytC (S189, E310 and H341) and the S15 of PorA that becomes mycoylated are labeled. Inset: active site view of the superposition of the MytC-acyl structure (salmon) and the AF model of MytC-PorA (same colors as in left)

## Discussion

Mycoloylation of porins in *C. glutamicum* has been discovered in 2010 (*14*) and later proved to be dependent, *in vivo,* on *mytC* expression (*19*). Here we set up an *in vitro* test showing that purified MytC is indeed able to mycoloylate PorA provided TMM is added to the reaction mixture. Importantly, another main mycoloyltransferase in *C. glutamicum*, MytA, is not able to mycoloylate PorA in identical experimental conditions. This result definitely proves that MytC is in fact the mycoloyltransferase directly involved in this atypical post-translational modification. It also indicates that MytC is able to recognize its acceptor substrate without the assistance of any other proteins although we cannot exclude that the involvement of other partners may improve the efficiency of the process. By using different synthetic analogs of TMM, we explored the specificity of MytC towards its mycolate donor and more particularly probed the importance of the mycolic acid motif. Our results clearly showed that the presence of the β–hydroxyl group of the meromycolate chain, a hallmark of all mycolic acids, is a key determinant of MytC specificity while the presence of the α-chain is not mandatory. This finding is particularly interesting if compared to MytA which, in contrast, does not have this strict specificity. Indeed, MytA is perfectly able to catalyze *in vitro* the transfer of a palmitoyl group from analog **6** (TMP) onto another TMP to form di-*O*-palmytoylated trehalose (an acyl analog of TDM) (Figure 4). MytA and MytC are therefore distinct mycoloyltransferases that evolved in their specificity both for donor and acceptor substrates. Interestingly, all these *in vitro* observations are consistent and may sustain *in vivo* data of the literature showing that deletion of *mytA*, but not of *mytC*, has a very dramatic effect on TDM synthesis (*7*). Conversely, deletion of *mytC* has been described to completely abolish PorA mycoloylation which can only be restored by ectopic expression of *mytC* but not of *mytA* (*19*). Concerning the promiscuity of MytA towards its mycolate donor, it is important to notice that acyl-TMM molecules have been detected in *C. glutamicum* according to two independent lipidomic studies (*34*, *35*). In the light of our results, it is therefore tempting to speculate that MytA could be responsible *in vivo* for their synthesis from acyl-trehalose molecules. In contrast, the strict specificity of MytC towards its mycolate donor seems essential to clearly distinguish protein mycoloylation from the *bona fide* Lgt/Lnt protein acylation machinery. The coexistence of two distinct lipidation pathways in *C. glutamicum* may suggest that each of them is dedicated to two independent subsets of proteins that may require distinct modification to adopt their active conformation or to be sorted to their final destination. In this context, it is tempting to speculate that MytC has evolved in order to avoid any unfortunate acylation of proteins that otherwise would lead to their mis-localization and/or mis-folding and may be deleterious for the cell. Finally, our finding that MytC, but not MytA, is very specific for β–hydroxylated acyl chains somehow support the choice of acyl-trehalose probes that have been developed recently in order to label the mycomembrane (*36*) but not those intended to identify the mycoloylome by metabolic labeling (*37*).

The MytC apoform structure (*19*) revealed that its catalytic triad adopted an inactive conformation and that its active site was blocked by a small helix. The ensemble of MytC structures and models highlights the highly flexible nature of its active site region, capable of adopting different conformations of the catalytic machinery. Upon formation of the acyl-intermediate, the blocking helix unfolded and was displaced from the active site pocket, providing space for the covalently bound acyl moiety. The H-bonding network of the catalytic triad however remained disrupted in the acyl-form. We hypothesize that this inactive form of the MytC-acyl enzyme is a temporary conformational that occurs before arrival of its acceptor substrate PorA. This probably represents a strategy to prevent futile hydrolysis of the reaction intermediate, as has been suggested for other mycoloyltransferases (*38*). The considerable flexibility of the MytC active site region is further illustrated by the restructuring of the residues 280 to 305, who transit from a disordered state in apo-MytC to an elongation of the α11 helix in the MytC-acyl enzyme. The towering position of this helix next to the active site pocket suggests that the α11 helix could be involved in interaction with the PorA substrate. Alphafold modeling of the MytC-PorA complex endorses this hypothesis. In this model the PorA protein forms a very amphiphilic helical bundle whose hydrophobic surface wraps around a hydrophobic patch of the helix α11 of MytC. The putative MytC binding site for PorA (residues 300 to 311) is disordered in the apo-form of the enzyme, but elongates the α11-helix in the structure of the acyl-intermediate. The PorA binding site overlaps with the regions of MytA and Ag85C involved in binding of the acceptor TMM to synthesize TDM (Supplementary Figure 8), suggesting that this region has evolved in MytC to accommodate PorA instead of TMM. Superposition of the MytC-PorA complex onto MytA, shows that 2 of the PorA helices overlap with 2 helices at the *N*-terminus of MytA that also blocked the active site region of the latter enzyme. Superposition of MytC-PorA onto the acyl-MytC also shows that the β-hydroxyl group could interact with PorA (inset Figure 6), explaining why this group might be a discriminating determinant for PorA mycoylation. The model also convincingly positions the serine 15 mycoloylation target facing the carbonyl of mycolate C13:0 bound to MytC.

Previous studies on mycoloyltransferases have provided important structural data but none of these included compounds that had the exact pattern of natural mycolic acids. For instance, a structure of the mycoloyltransferase Ag85C-acyl-enzyme was obtained in an elegant way using THL, a versatile lipid esterase covalent inhibitor. Indeed, this natural compound features two chiral carbons that mimic the core attributes of the mycolic acid. The 2-alkyl, 3-hydroxy fatty acid moiety has however an inverted configuration with respect to the asymmetric carbons of the natural mycolic acids. While the two THL carbons of interest have (2*S*, 3*S*) configurations, the natural mycolic acids have an opposite (*R*) configuration at the two chiral carbons, C2 and C3 (*39*). Furthermore, THL carries a voluminous peptidyl side arm which is absent in mycolic acids. The structure of the Ag85C-acyl-enzyme, combined with molecular dynamics, provided interesting hypothesis regarding the mechanism of the transesterification reaction: *i*) the binding of the incoming acceptor molecule could drive Ag85C to form an active conformation and *ii*) the β-hydroxy of MA would directly or indirectly activate the incoming nucleophile. When comparing MytC acylated with a C13:0 mycolate carrying the native carbon configurations with the Ag85C-acyl-enzyme, we noticed that their ester moieties bound to the catalytic serine are in opposite orientations (Supplementary Figure 5). This implies a significantly different positioning of the electrophilic carbonyl to be attacked by the incoming nucleophilic alcohol of the acceptor. Interestingly, despite the opposite configuration of the C3, the β-hydroxyl group is accommodated in a relatively similar way. Consequently, the two alkyl chains do not project in the same direction in the two structures. Here, the α-chain is perfectly positioned in the hydrophobic pocket, while the meromycolic chain remains in another more interfacial cleft, directed towards the exit of the active site cave. This is a clear difference with the previously proposed model based on the Ag85C-THL structure (*40*) and that does not necessarily exclude an interfacial mechanism “scooting mechanism” as proposed previously.

We finally extract from our biochemical and structural studies a hypothetical mechanism of the MytC catalyzed reaction (Supplementary Figure 10). In the resting state the active site pocket of the apo-enzyme is locked by a small helix (η3) which unfolds upon entrance of the TMM mycolate donor into the active site. After formation of the intermediate ester, the crucial β-hydroxyl group could either interact with the catalytic triad or activate the carbonyl for nucleophilic attack through intramolecular hydrogen-bond formation. The inactive configuration of the catalytic triad protects the intermediate from being hydrolyzed. Binding of the PorA mycolate acceptor restores the active configuration of the catalytic triad and through interactions with the α8 and α11 helices of MytC inserts its serine 15 in the catalytic site for the subsequent transesterification.

## Conclusion

Lipoproteins are essential in bacteria where they represent between 2 to 3% of total proteins and are involved in various functions such as cell physiology, nutrients influx, drug efflux or pathogenicity. Whereas in *Corynebacteriales*, acylated proteins have been well characterized, the protein *O*-mycoloylation has only recently been discovered in *C. glutamicum* as a new type of protein lipidation. It could be a central process in cell envelope organization and function and also represent an interesting therapeutic target. In this study, the MytC-mycolate acyl-enzyme structure and the structure-activity relationship data obtained from a collection of synthetic TMM analogues clearly show a unique mechanism and specificity within the mycoloyltransferases family. In particular, the role of the β-hydroxyl group of mycolic acid, which had previously been suggested as important in the Ag85C mechanism, is here unambiguously demonstrated as the key determinant to specifically individualize acylation and mycoloylation pathways in the cell. We also observed the presence of an efficient enzyme locking system to avoid hydrolysis processes in this transesterification reaction. Altogether, these findings provide additional knowledges in our understanding of the specificity and mechanism of the mycoloyltransferases whose apparent redundancy is still not completely understood in *Corynebacteriales*. These results could be useful in designing specific inhibitors for each individual mycoloyltransferases which could be interesting targets for innovative therapeutic approaches.

## Materials and Methods

### Bacterial Strains and Growth Conditions

All *C. glutamicum* strains used in this study are derivative of the strain ATCC 13032 RES167. *C. glutamicum* ATCC13032 RES167 was grown in brain heart infusion (BHI) medium with shaking (220 rpm) at 30°C.

### Plasmids Construction

The plasmids used in this study for protein expression were constructed previously. MytA_his_ encoding gene was cloned in pCGL482 under its own promoter and signal sequence (*41*) and introduced in *C. glutamicum ΔmytA* strain. MytC_his_ encoding gene was cloned in pCGL482 under its own promoter and signal sequence (*19*) and introduced in *C. glutamicum* ATCC 13032 Δ*mytC* strain. MytC_his_ is highly expressed (5 mg.L^-1^) and partly secreted in the culture supernatant or localized in the cell envelope. PorA_his_ encoding gene was cloned in pXMJ19 under a *tac* promoter (*16*) and introduced in *C. glutamicum* ATCC 13032 Δ*mytC* strain. PorA_his_ is moderately expressed and essentially found in the cell envelope.

### Expression and Purification of PorA_his_

Recombinant PorA protein was purified either from WT or Δ*mytC* strain according to Issa et al. (*17*). Proteins were dialyzed in Tris-HCl 10 mM, NaCl 10 mM, pH 8.0 buffer and aliquots were stored at −20°C. The purity and homogeneity of proteins were analyzed on 16% Tricine SDS-PAGE (*42*).

### Expression and Purification of MytC_his_ and MytA_his_

#### Purification of MytA_his_

For large scale purification of MytA_his_, cells (*C. glutamicum* Δ*mytA* strain transformed with *p*CGL482-*mytAhis*) from 1.2 L overnight cultures (BHI + Cam 6 µg/ml, 30°C) were removed by centrifugation at 7000 x g at 4°C. Proteins were then precipitated from the supernatant by adding ammonium sulfate to 70% saturation. Incubation was carried out for 1h at 4°C with constant shaking. After centrifugation at 7000 x g for 15 min at 4°C, the protein-containing pellet was resuspended in 60 mL of Tris 25 mM buffer pH 7.5, NaCl 200 mM (purification buffer). The solution was dialyzed for 24h at 4°C against the purification buffer (Spectrum, Spectra/Por MWCO 10 kDa) before loading on a 1 mL Ni-nitrilotriacetic acid (NTA) column. The flow through was recovered and loaded one more time on the column. Proteins weakly associated with the Ni-NTA-resin were washed off by running 40 mL of purification buffer. Elution was finally performed with 3 column volumes of purification buffer containing imidazole at 250 mM. Fractions containing MytA_his_ were dialyzed in 10 mM phosphate buffer pH 8.0 and 10 mM NaCl. The purification was controlled by running 12% SDS PAGE (see Supplementary Figure 1a) gels after ammonium sulfate, Ni-NTA, and dialysis. The concentration of MytA_his_ (χ_280_ = 140,385 M^-1^cm^-1^, 66,982 Da) was determined using absorbance at 280 nm.

#### Purification of MytC_his_

For large-scale purification, Δ*mytC* (*MytC_his_*) cells were recovered after overnight culture by centrifugation at 6,000 rpm at 10°C. Proteins were then precipitated from the solution by adding ammonium sulfate to 70% saturation. Incubation was carried out for 1h at room temperature with constant shaking. After centrifugation at 6,000 rpm for 15 min at 10°C, the protein-containing pellet was resuspended in 60 mL of 25 mM Tris, 200 mM NaCl (pH 7.5) (purification buffer). The solution was dialyzed for 24h at 4°C against the purification buffer (Spectra/Por, molecular mass cutoff of 10 kDa; Spectrum), and then mixed with the Ni-(NTA) resin. After 2h incubation at 4°C and a centrifugation at 2,000 rpm for 10 min at 4°C, the Ni-NTA resin was recovered. Proteins weakly associated with the Ni-NTA resin were washed off by adding 20 mL of purification buffer. After 2 washes, elution was finally performed with 4 resin volumes of purification buffer containing imidazole at 250 mM. Fractions 1 to 4 were then dialyzed in 500 mL of 10 mM Tris, 10 mM NaCl (pH 8.0) (reaction buffer). The whole purification procedure (Ni-NTA) was always done on the same day, and aliquots of CgMytC-His were conserved at −80C. The purification was controlled by 12% SDS-PAGE gels (see Supplementary Figure 1a) after ammonium sulfate and Ni-NTA steps. Protein quantification was performed by using a Nanodrop spectrophotometer after the final dialysis (χ_280_ = 86,525 M^-1^cm^-1^, 37,451 Da).

### PorA mycoloylation assay

The protein mycoloylation activity of MytC was evaluated by using non-mycoloylated PorA_his_ as a substrate. PorA_his_ mycoloylation was assessed by SDS PAGE and MALDI TOF analysis. A standard reaction mixture (30 µL) consisting of 6 μM PorA_his_, 5 μM MytC, 1 mM of mycolate donor (TDM/TMM mix, synthetic TMM or TMM analogs) was incubated at 37°C for 3h in reaction buffer (Tris-HCl 10 mM, NaCl 10 mM, pH 8.0). The reaction was stopped either by adding loading buffer (30 µL) before electrophoresis analysis on SDS Tricine gels or, by adding the matrix for MALDI-TOF analysis (sinapinic acid, 20 mg/mL in H_2_O/CH_3_CN 1:1).

### Trehalose mycoloylation assay

A standard reaction mixture (50 µL) consisting of 5 μM MytC or MytA, 1 mM of sTMM was incubated at 37°C for 3h in reaction buffer (Tris-HCl 10 mM NaCl 10 mM, pH 8.0). The reaction was stopped at −20°C. Aliquots were then mixed with the matrix for MALDI-TOF analysis (6-aza-2-thiothymine,10 mg/mL in H_2_O/CH_3_CN 1/1) or loaded on TLC and SDS-PAGE. Similar reactions were run by using TMM C13:0 and TMP instead of sTMM at 1 mM final concentration.

### Preparation of TDM/TMM mix and purification of TMM

Trehalose mycolates were purified from membrane vesicles secreted by the strain 13032 Δ*aftB* (*43*). Bacteria were grown in BHI (600 mL) overnight and the culture supernatant was recovered after two successive centrifugations (6000 g, 15 min 4°C). Vesicles were isolated by ultracentrifugation (35,000 rpm for 2h at 4°C in a 45 Ti rotor) and resuspended in 20 mL of Hepes 25 mM pH 7.4 at 4°C and membrane lipids extracted by addition of 20 mL chloroform and 40 mL methanol. After 2h incubation, 20 mL of chloroform and 20 mL of H_2_0 were added and the solution further incubated for 24h. The chloroform phase was recovered and evaporated. Lipids were resuspended in 2 mL and stored at −20°C. At this stage the lipid extract (TDM/TMM mix) is mainly enriched in TMM and TDM as shown in Supplementary Figure 3.

For TMM purification, the TDM/TMM mix was subjected to a preparative thin layer chromatography on a silica plate using a solvent mixture composed of chloroform/methanol/H_2_0 34:15:2. TMM was recovered by scratching the plate at the dedicated migration position of the lipid by comparison with a reference plate migrating simultaneously and revealed by vaporating a mixture of ethanol and sulfuric acid (90:10) on the plate which is then heated at 130°C until appearance of the bands (see Supplementary Figure 1b).

### Lipid analysis by thin layer chromatography

Lipids were analyzed by TLC using the standard procedure already described by Dietrich *et al*.(*41*)

### Mass spectrometry MALDI-TOF analysis

MALDI-TOF spectra were acquired on an Axima performance mass spectrometer (Shimadzu corporation) equipped with a pulsed nitrogen laser emitting at 337 nm and an accelerating voltage of 20 kV. All spectra were acquired in the positive linear or reflectron mode.

For MytC *in vitro* assay, a mixture of 1 µL of sample and 1 µL of sinapinic acid matrix solution (20 mg/mL in H_2_O:CH_3_CN 1:1, TFA 0.1%) was deposited onto the MALDI plate and allowed to dry in air. External mass calibration was performed thanks to SpheriCal Aqua Protein Low kit (Sigma).

For the trehalose mycoloylation *in vitro* assay a mixture of 1 µL of sample and 1 µL of 6-aza-2-thiothymine (ATT) matrix solution (10 mg/mL in H_2_O:CH_3_CN 1:1) were deposited onto the MALDI plate and allowed to air dry. External calibration was performed with pepmix XT kit (Fisher Scientific).

### TMM analogs synthesis

Synthetic details and characterization of all previously unreported compounds are provided in supplementary materials.

### MytC crystallization with TMM analogs and resolution of the protein structures

The MytC (126.5 µM) and TMM C13:0 or TMP (1.265mM) mixtures were incubated at room temperature for 1h and then co-crystallized by sitting-drop vapor diffusion at 18°C (291°K) from a 0.1 nanoL:0.1 nanoL mixture of protein complex solution with crystallization solutions. Crystals of the MytC-TMM C13:0 complex were obtained using a crystallization solution composed of 0.1M sodium chloride, 1.4 M ammonium sulfate, 0.1 M Hepes pH 7.1. The MytC-TMM C13:0 crystals were transferred to a solution composed of 1.5 M ammonium sulfate, 0.1 M sodium chloride, 0.1 M Hepes pH 7.1 and 23% of glycerol incremented with 0.5 mM TMM-C13:0 before flash-cooled in liquid nitrogen. The crystallization solution for the MytC-TMP crystals contained 0.2 M magnesium chloride, 0.1 M TRIS pH 8.5 and 32 %(w/v) PEG 4000. For diffraction experiments the MytC-TMP crystals were transferred to the crystallisation solution incremented with 15% Glycerol and 0.5mM TMP before flash-cooled in liquid nitrogen. Diffraction data were recorded on beam line Proxima2 (synchrotron SOLEIL, France) and were processed using the XDS package (*44*). The structure was determined by the molecular replacement method using the structure of apo-MytC as model (PDB accession code 4H18) and the program Molrep (*45*) implemented in ccp4 (*46*). The model was further improved by iterative cycles of manual rebuilding using COOT (*47*) and refinement using BUSTER program (Bricogne 2017, BUSTER version 2.10.3. Cambridge, United Kingdom: Global Phasing Ltd.). Statistics for data collection and refinement are summarized in Supplementary Table 1. The atomic coordinates (and structure factors) or the MytC-TMM C13:0 complex have been deposited into the Brookhaven Protein Data Bank under the accession number 8QHF.

### Alphafold models

The models were obtained using the web site: https://colab.research.google.com/github/sokrypton/ColabFold/blob/main/beta

(see the input data parameters in supplementary Table 1)

## Supporting information

Supplementary information Lesur et al.

## Acknowledgments

This work benefited from the expertise of the crystallization platform of I2BC, supported by the French Infrastructure for Integrated Structural Biology (FRISBI, ANR-10-INSB-05-05). We acknowledge the synchrotrons ESRF (Grenoble, France) and SOLEIL (Saint-Aubin, France) for provision of synchrotron radiation facilities and we would like to thank the staffs of beamlines PROXIMA-2A and PROXIMA-1 at SOLEIL, and 1D23-2 at ESRF for assistance and advices during data collection. We are very grateful to Mohamed Chami for his very useful advices concerning the preparation and solubilization of mycolate lipids. We are grateful to all members of the lab for their support and advices. We thank the Ministère de l’enseignement supérieur et de la recherche for grants to EL and PR, the China Scolarship Council (CSC grant N° 202006230063 for YZ). We also thank the Agence Nationale de la Recherche (PTMyco, grant NO. ANR-22-CE44-0005-03).

## Contributions

ND, YB, DG and NB designed the strategy and the experiments. EL, YZ, ND, NL, ILSG, CD, PR, DU, GD and LA made the experiments. YB, DG, ILSG, HVT, FCB, and NB wrote the manuscript.

## Bibliography

1. H. Marrakchi, M.-A. Lanéelle, M. Daffé, Chem. Biol. 21, 67–85 (2014).

2. D. Portevin et al., Proc. Natl. Acad. Sci. U. S. A. 101, 314–319 (2004).

3. S. Gavalda et al., Chem. Biol. 21, 1660–1669 (2014).

4. J. Li et al., Cell. 176, 636–648.e13 (2019).

5. M. Daffé, H. Marrakchi, Microbiol. Spectr. 7 (2019).

6. N. Dautin et al., Biochim. Biophys. Acta - Gen. Subj. 1861, 3581–3592 (2017).

7. S. Brand, K. Niehaus, A. Pühler, J. Kalinowski, Arch. Microbiol. 180, 33–44 (2003).

8. C. De Sousa-D’Auria et al., FEMS Microbiol. Lett. 224, 35–44 (2003).

9. M. Faller, M. Niederweis, G. E. Schulz, Science. 303, 1189–1192 (2004).

10. Q. Wang et al., Science. 367, 1147–1151 (2020).

11. M. Niederweis, E. Maier, T. Lichtinger, R. Benz, R. Kramer, J. Bacteriol. 177, 5716– 5718 (1995).

12. T. Lichtinger, A. Burkovski, M. Niederweis, R. Krämer, R. Benz, Biochemistry. 37, 15024–15032 (1998).

13. K. Ziegler, R. Benz, G. E. Schulz, J. Mol. Biol. 379, 482–491 (2008).

14. E. Huc et al., J. Biol. Chem. 285, 21908–21912 (2010).

15. E. Huc et al., J. Bacteriol. 195, 4121–8 (2013).

16. P. Rath et al., J. Biol. Chem. 286, 32525–32 (2011).

17. H. Issa et al., PLoS One. 12 (2017), doi:10.1371/journal.pone.0171955.

18. C. Carel et al., Proc. Natl. Acad. Sci. U. S. A. 114, 4231–4236 (2017).

19. E. Huc et al., J. Bacteriol. 195 (2013).

20. F. Migliardo et al., Chem. Phys. Lipids. 223 (2019).

21. P. Rath et al., Biochim. Biophys. Acta - Biomembr. 1828, 2173–2181 (2013).

22. E. Lesur, P. Rollando, D. Guianvarch, Y. Bourdreux, Comptes rendus. Chim. Online first (2023); pp. 1–22. doi 10.5802/crchim.246. (2023).

23. E. Lesur et al., Chem. Commun. 55 (2019), doi:10.1039/c9cc05754d.

24. R. Noyori et al., J. Am. Chem. Soc. 109, 5856–5858 (1987).

25. G. Fráter, Helv. Chim. Acta. 62, 2825–2828 (1979).

26. D. Seebach, D. Wasmuth, Helv. Chim. Acta. 63, 197–200 (1980).

27. C. De Sousa-D’Auria et al., FEMS Microbiol. Lett. 224, 35–44 (2003).

28. J. T. Belisle et al., Science. 276, 1420–2 (1997).

29. J. Boucau, A. K. Sanki, B. J. Voss, S. J. Sucheck, D. R. Ronning, Anal. Biochem. 385, 120–7 (2009).

30. L. Favrot et al., Nat. Commun. 4, 2748 (2013).

31. D. R. Ronning et al., Nat. Struct. Biol. 7, 141–6 (2000).

32. D. R. Ronning, V. Vissa, G. S. Besra, J. T. Belisle, J. C. Sacchettini, J. Biol. Chem. 279, 36771–7 (2004).

33. C. M. Goins, S. Dajnowicz, M. D. Smith, J. M. Parks, D. R. Ronning, J. Biol. Chem. 293, 3651–3662 (2018).

34. S. Klatt et al., J. Lipid Res. 59, 1190–1204 (2018).

35. H. Y. J. Wang, R. V. V. Tatituri, N. K. Goldner, G. Dantas, F. F. Hsu, Biochimie. 178, 158–169 (2020).

36. H. L. Hodges, R. A. Brown, J. A. Crooks, D. B. Weibel, L. L. Kiessling, Proc. Natl. Acad. Sci. U. S. A. 115, 5271–5276 (2018).

37. H. W. Kavunja et al., Chem. Commun. (Camb). 52, 13795–13798 (2016).

38. K. M. Backus et al., J. Biol. Chem. 289, 25041–53 (2014).

39. C. Asselineau, G. Tocanne, J. F. Tocanne, Bull. Soc. Chim. Fr. 4, 1445–1449 (1970).

40. C. M. Goins, C. M. Schreidah, S. Dajnowicz, D. R. Ronning, J. Biol. Chem. 293, 1363–1372 (2018).

41. C. Dietrich et al., Mol. Microbiol. (2020).

42. H. Schägger, Nat. Protoc. 1, 16–22 (2006).

43. R. Bou Raad et al., J. Bacteriol. 192, 2691–2700 (2010).

44. W. Kabsch, Acta Crystallogr. Sect. D Biol. Crystallogr. 66, 125–132 (2010).

45. A. Vagin, A. Teplyakov, Acta Crystallogr. Sect. D Biol. Crystallogr. 66, 22–25 (2010).

46. J. Agirre et al., Acta Crystallogr. Sect. D, Struct. Biol. 79, 449–461 (2023).

47. P. Emsley, K. Cowtan, Acta Crystallogr. Sect. D Biol. Crystallogr. 60, 2126–2132 (2004).

