## Supplementary information Lesur et al. for "Synthetic mycolates derivatives as molecular tools to decipher protein mycoloylation, a unique post-translational modification in bacteria"

Affiliations:

#### 1- Chemical Synthesis

**General methods:**

All air sensitive reactions were carried out in oven-dried glassware under a slight positive pressure of argon. Solvents were dried by standard methods. THF was distilled from sodium benzophenone ketyl. TLC (Silica Gel 60 $F_{254}$) were visualized under UV (254 nm) and by staining either in 5% ethanolic sulfuric acid or orcinol or phosphomolybdic acid. Silica gel SDS 60 ACC 35-70 µm was used for column chromatography. NMR spectra were recorded on Bruker DRX 300 or AV 360 spectrometers. Chemical shifts (in ppm) were determined relative to residual undeuterated solvent as an internal reference. Abbreviations of multiplicity were as follows: s (singlet), d (doublet), dd (doublet of doublet), t (triplet), at (apparent triplet), m (multiplet), b (broad). Coupling constants in hertz (Hz) were measured from one-dimensional spectra. High-resolution mass spectra (positive or negative mode ESI) were performed on a Bruker Daltonics micrOTOF-QII spectrometer. Optical rotations were measured on an Anton Paar MCP 150 polarimeter (*c* in g / 100 mL). Synthetic TMM, TDM, **9b** and **10b** were synthesized according to the literature.[i,ii,iii]

**2-Tetradecyl-octadecanoic acid 8**

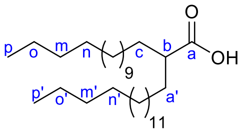


To a solution of dry diisopropylamine (830 µL, 5.92 mmol, 2.5 equiv) in THF (5 mL) cooled to 0 °C was added dropwise *n*-butyllithium (3.5 mL, 1.6 M, 5.60 mmol, 2.4 equiv). After stirring for 30 min at 0 °C the solution was added, *via* cannula, to a solution of commercially available palmitic acid (600 mg, 2.34 mmol) in dry THF (10 mL). The reaction mixture was stirred for 1 h at 0 °C and then, iodohexadecane (800 µL, 2.76 mmol, 1.2 equiv) and HMPA (500 µL, 2.87 mmol, 1.2 equiv) were added slowly. The mixture was warmed to 50 °C and stirred overnight at this temperature. The mixture was acidified with an aqueous HCl solution (1M, pH ≈ 1) and extracted three times with EtOAc. The combined organic layers were washed with an aqueous NaCl solution, dried over $Na_2SO_4$ and concentrated under reduced pressure. The residue was purified by flash silica gel chromatography (cyclohexane/EtOAc, 13:1 to 12:1) to give compound **8** (515 mg, 46%) as a colourless oil. NMR data were in agreement with literature.[iv] $R_f$ = 0.45 (cyclohexane/EtOAc, 13:1). $^1$H NMR ($CDCl_3$, 360 MHz) δ (ppm): 2.35 (m, 1H, $H_b$), 1.70-1.39 (m, 4H, $H_c$ and $H_{a'}$), 1.40-1.12 (m, 52H, $H_c$-$H_o$ and $H_{b'}$-$H_{o'}$), 0.90 (t, 6H, $J$ = 7.0 Hz, $H_{p'}$ and $H_p$). $^{13}$C NMR ($CDCl_3$, 90 MHz) δ (ppm): 180.2 ($C_a$), 45.4 ($C_b$), 32.1, 31.9, 29.6, 29.5, 29.4, 29.3, 27.3, 22.6 (28C, $C_c$-$C_o$, $C_{a'}$-$C_{o'}$), 14.1 (2C, $C_p$, $C_{p'}$). HRMS (ESI): calcd for $C_{32}H_{63}O_2$ $[M-H]^-$ 479.4834, found 479.4812.

**Methyl 3-oxo-octanoate 9a:**

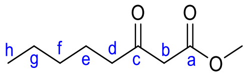


Compound **9a** was obtained according to Masamune's procedure.[v] Briefly, carbonyldiimidazole (1.68 g, 10.32 mmol, 1.2 equiv) was added to a solution of commercially available hexanoic acid (1.1 mL, 8.80 mmol) in dry THF (84 mL). After stirring at room temperature for 6-8 h, the magnesium salt of monomethylmalonate (2.89 g, 11.22 mmol, 1.3

equiv) was added. The reaction mixture was stirred for 18 h at room temperature and was then acidified with an aqueous HCl solution (1M, pH ≈ 1). The reaction mixture was extracted three times with EtOAc, and the combined organic layers were dried over $Na_2SO_4$ and concentrated under reduced pressure. The residue was purified by flash silica gel chromatography (cyclohexane to cyclohexane/EtOAc 99:1) to give compound **9a** (869 mg, 59%) as a colourless oil. Spectral data were in agreement with literature.[vi] $R_f$ = 0.49 (cyclohexane/EtOAc, 85:15). $^1$H NMR ($CDCl_3$, 300 MHz) δ (ppm): 3.74 (s, 3H, $OCH_3$), 3.44 (s, 2H, $H_b$), 2.52 (t, 2H, $J$ = 7.4 Hz, $H_d$), 1.66-1.52 (m, 2H, $H_e$), 1.34-1.22 (m, 4H, $H_f$, $H_g$), 0.89 (t, 3H, $J$ = 6.9 Hz, $H_h$). $^{13}$C NMR ($CDCl_3$, 75 MHz) δ (ppm): 203.0 ($C_c$), 167.8 ($C_a$), 52.5 ($OCH_3$), 49.2 ($C_b$), 43.2 ($C_d$), 31.3 ($C_f$), 23.3 ($C_e$), 22.5 ($C_g$), 14.0 ($C_h$). HRMS (ESI): calcd for $C_9H_{16}NaO_3$ $[M+Na]^+$ 195.0992, found 195.0983, calcd for $C_9H_{17}O_3$ $[M+H]^+$ 173.1172, found 173.1166.

**Methyl 3-oxo-tridecanoate 9c**

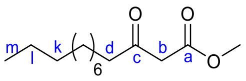


Compound **9c** was synthetized according to Masamune's procedure.[v] Carbonyldiimidazole (1.04 g, 6.44 mmol, 1.2 equiv) was added to a solution of commercially available undecanoic acid (1 g, 5.37 mmol) in dry THF (55 mL). After stirring at room temperature for 6-8 h, the magnesium salt of monomethylmalonate (1.80 g, 6.98 mmol, 1.3 equiv) was added. The reaction mixture was stirred for 18 h at room temperature and was then acidified with an aqueous HCl solution (1M, pH ≈ 1). The reaction mixture was extracted three times with EtOAc, and the combined organic layers were dried over $Na_2SO_4$ and concentrated under reduced pressure. The residue was purified by flash silica gel chromatography (cyclohexane/EtOAc, 98:2 to 95:5) to give compound **9c** (679 mg, 52%) as a colourless oil. NMR data were in agreement with literature.[vii] $R_f$ = 0.50 (cyclohexane/EtOAc, 9:1). $^1$H NMR ($CDCl_3$, 300 MHz) δ (ppm): 3.74 (s, 3H, $OCH_3$), 3.45 (s, 2H, $H_b$), 2.52 (t, 2H, $J$ = 7.5 Hz, $H_d$), 1.65-1.54 (m, 2H, $H_e$), 1.34-1.20 (m, 14H, $H_f$-$H_l$), 0.87 (t, 3H, $H_m$). $^{13}$C NMR ($CDCl_3$, 75 MHz) δ (ppm): 203.1 ($C_c$), 167.9 ($C_a$), 52.5 ($OCH_3$), 49.2 ($C_b$), 43.2 ($C_d$), 32.0, 29.7, 29.6, 29.5, 29.4, 29.1, 22.8 (7C, $C_f$-$C_l$), 19.8 ($C_k$), 23.6 ($C_e$), 14.3 ($C_m$). HRMS (ESI): calcd for $C_{14}H_{26}NaO_3$ $[M+Na]^+$ 265.1774, found 265.1766, calcd for $C_{14}H_{27}O_3$ $[M+H]^+$ 243.1955, found 243.1943.

**Methyl (*R*)-3-hydroxy-octanoate 10a**

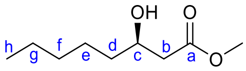


The (*R*)-BINAP-$RuBr_2$ complex was prepared under argon according to a reported procedure.[viii] To a solution of (*R*)-BINAP (77 mg, 0.12 mmol, 0.022 equiv) and (Cod)Ru(2-methylallyl)$_2$ (36 mg, 0.11 mmol, 0.02 equiv) in anhydrous and degassed acetone (6 mL), were added 1.5 mL of a methanolic HBr solution (0.165 M, 0.25 mmol, 0.044 equiv). The reaction mixture was stirred for 30 min at room temperature and then concentrated under vacuum. A solution of β-ketoester **9a** (970 mg, 5.6 mmol) in degassed and dry MeOH (6.9 mL) was added *via* cannula to the catalyst. The mixture was purged three times with dihydrogen and vigorously stirred overnight at 50 °C under 1 atm of dihydrogen. The solvent was evaporated under vacuum and the residue was purified by silica gel chromatography (cyclohexane/EtOAc, 99:1 to 95:5) to give the desired product **10a** (878 mg, 90%) as a colourless oil. NMR data were in agreement with literature.[ix] $R_f$ = 0.23 (cyclohexane/EtOAc, 85:15). $^1$H NMR ($CDCl_3$, 360 MHz) δ (ppm): 4.00 (m, 1H, $H_c$), 3.71 (s, 3H, $OCH_3$), 2.52 (dd, 1H, *J* = 16.4 Hz, *J* = 3.2 Hz, $H_b$), 2.41 (dd, 1H, *J* = 16.4 Hz, *J* = 8.7 Hz, $H_{b'}$), 1.59-1.23 (m, 8H, $H_d$-$H_g$), 0.89 (t, 3H, *J* = 6.5 Hz, $H_h$). $^{13}$C NMR ($CDCl_3$, 90 MHz) δ (ppm): 173.4 ($C_a$), 68.0 ($C_c$), 51.7 ($OCH_3$), 41.3 ($C_b$), 36.6, 31.7, 25.2, 22.6 (4C, $C_d$-$C_g$), 14.0 ($C_h$). HRMS (ESI): calcd for $C_9H_{18}NaO_3$ $[M+Na]^+$ 197.1148, found 197.1143. $[\alpha]_D^{20}$ = -19 (*c* 1.07, $CHCl_3$).

**Methyl (*R*)-3-hydroxy-tridecanoate 10c**

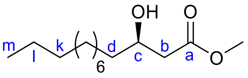


The (*R*)-BINAP-$RuBr_2$ complex was prepared under argon according to a reported procedure.[viii] To a solution of (*R*)-BINAP (41.9 mg, 0.067 mmol, 0.022 equiv) and (Cod)Ru(2-methylallyl)$_2$ (19.5 mg, 0.061 mmol, 0.020 equiv) in anhydrous and degassed acetone, were added 820 µL of a methanolic HBr solution (0.165 M, 0.135 mmol, 0.044 equiv). The reaction mixture was stirred for 30 min at room temperature and then concentrated under vacuum. A solution of β-ketoester **9c** (742 mg, 3.06 mmol) in degassed and dry MeOH (3.75 mL) was added *via* cannula to the catalyst. The mixture was purged three times with dihydrogen and vigorously stirred overnight at 50 °C under 1 atm of dihydrogen. The solvent was evaporated under vacuum and the residue was purified by silica gel chromatography (cyclohexane/EtOAc, 95:5 to 90:10) to give the desired product **10c** (570.1 mg, 76%) as a colourless oil. $R_f$ = 0.21 (cyclohexane/EtOAc, 9:1). $^1$H NMR ($CDCl_3$, 360 MHz) δ (ppm): 3.97 (m, 1H, $H_c$), 3.69 (s, 3H, $OCH_3$), 2.95 (s, 1H, OH), 2.49 (dd, 1H, *J* = 16.3 Hz, *J* = 3.2 Hz, $H_b$), 2.38 (dd, 1H, *J* = 16.3 Hz, *J* = 8.9 Hz, $H_{b'}$), 1.54-1.16 (m, 18H, $H_d$-$H_l$), 0.85 (t, 3H, $H_m$). $^{13}$C NMR ($CDCl_3$, 90 MHz) δ (ppm): 173.6 ($C_a$), 68.1 ($C_c$), 51.8 ($OCH_3$), 41.2 ($C_b$), 36.6 ($C_d$), 32.0, 29.7, 29.6, 29.4, 25.6, 22.8 (9C, $C_d$-$C_l$), 14.2 ($C_m$). HRMS (ESI): calcd for $C_{14}H_{28}NaO_3$ $[M+Na]^+$ 267.1931, found 267.1924. $[\alpha]_D^{20}$ = -14 (*c* 1.13, $CHCl_3$).

**(*R*)-3-triethylsilyloxy-octadecanoic acid 11**

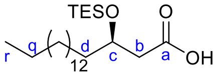


A 10% aqueous solution of NaOH (16 mL) was added to a solution of compound **10b**[iii] (501.9 mg, 1.60 mmol) in MeOH (16 mL). The reaction mixture was stirred overnight at room temperature and then acidified to pH ≈ 1 with an aqueous HCl solution (1M). The mixture was extracted with EtOAc, the combined organic layers were dried over $Na_2SO_4$ and concentrated under reduced pressure. The crude product was purified by flash silica gel chromatography (EtOAc/MeOH, 99:1 to 97:3) to give the corresponding carboxylic acid (426.3 mg, 89%) as a colourless syrup. NMR data were in agreement with literature.[iii] $R_f$ = 0.1 (cyclohexane/EtOAc, 1:1). $^1$H NMR ($CDCl_3$, 360 MHz) δ (ppm): 4.04 (m, 1H, $H_c$), 2.60 (dd, 1H, *J* = 16.7 Hz, *J* = 3.2 Hz, $H_b$), 2.49 (dd, 1H, *J* = 16.5 Hz, *J* = 8.9 Hz, $H_{b'}$), 1.65-1.15 (m, 28H, $H_d$-$H_q$), 0.89 (t, 3H, J = 7.0 Hz, $H_r$). $^{13}$C NMR ($CDCl_3$, 75 MHz) δ (ppm): 175.0 ($C_a$), 68.0 ($C_c$), 36.5 ($C_b$), 31.8, 29.5, 29.2, 25.4, 22.5 (14C, $C_d$-$C_q$), 13.8 ($C_r$). HRMS (ESI): calcd for $C_{18}H_{36}NaO_3$ $[M+Na]^+$ 323.2557, found 323.2551. $[\alpha]_D^{20}$ = -1 (*c* 1.0, $CHCl_3$/MeOH 1/1). To a solution of this compound (92 mg, 0.3 mmol) in dry pyridine (2 mL) was added dropwise TESCl (115 µL, 0.67

mmol, 2.2 equiv). The reaction mixture was stirred overnight at room temperature and was then diluted with a saturated aqueous NaCl solution. The reaction mixture was extracted three times with EtOAc. The combined organic layers were dried over $Na_2SO_4$ and concentrated under reduced pressure. The residue was purified by flash silica gel chromatography (cyclohexane/EtOAc, 80:20) to give compound **11** (75 mg, 59%) as a colourless oil. $R_f$ = 0.60 (cyclohexane/EtOAc 8:2). $^1$H NMR ($CDCl_3$, 360 MHz) δ (ppm): 4.05 (m, 1H, $H_c$), 2.60 (dd, 1H, *J* = 15.4 Hz, *J* = 5.0 Hz, $H_b$), 2.50 (dd, 1H, *J* = 15.4 Hz, *J* = 4.5 Hz, $H_{b'}$), 1.62-1.46 (m, 2H, $H_d$), 1.33-1.19 (m, 26H, $H_e$-$H_q$), 0.98 (t, 9H, *J* = 7.6 Hz, $Si(CH_2C\mathit{H}_3)_3$), 0.88 (t, 3H, *J* = 7.0 Hz, $H_r$), 0.66 (q, 6H, *J* = 7.6 Hz, $Si(C\mathit{H}_2CH_3)_3$). $^{13}$C NMR ($CDCl_3$, 90 MHz) δ (ppm): 174.6 ($C_a$), 69.6 ($C_c$), 41.6 ($C_b$), 37.3 ($C_d$), 32.0, 29.8, 29.7, 29.5, 25.5, 22.9 (13C, $C_e$-$C_q$), 14.3 ($C_r$), 6.9 ($Si(CH_2\mathit{C}H_3)_3$), 5.0 ($Si(\mathit{C}H_2CH_3)_3$). HRMS (ESI): calcd for $C_{24}H_{50}NaO_3Si$ $[M+Na]^+$ 437.3421, found 437.3400. $[\alpha]_D^{20}$ = - 1 (*c* 0.96, $CHCl_3$).

##### Methyl (2*R*, 3*R*)-2-pentyl-3-hydroxy-octanoate 13a

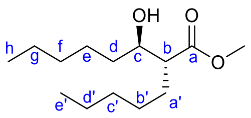


To a solution of dry diisopropylamine (2 mL, 14.27 mmol, 2.8 equiv) in THF (13.6 mL) cooled to -40 °C were added dropwise *n*-butyllithium (8 mL, 1.6 M, 12.8 mmol, 2.5 equiv). After stirring for 45 min at -40 °C, the mixture was cooled to -78 °C and a solution of β-hydroxyester **10a** (878 mg, 5.04 mmol) in dry THF (29 mL) was added *via* cannula. The reaction mixture was stirred for 1h30 at -78 °C then iodopentane (2 mL, 15.12 mmol, 3 equiv) and HMPA (1.05 mL, 6.05 mmol, 1.2 equiv) were added slowly. The mixture was warmed slowly to -10 °C and stirred overnight at this temperature. The mixture was diluted with a saturated aqueous $NH_4Cl$ solution and extracted three times with EtOAc. The combined organic layers were washed with an aqueous NaCl solution, dried over $Na_2SO_4$ and concentrated under reduced pressure. The residue was purified by flash silica gel chromatography (cyclohexane/EtOAc, 95:5) to give compound **13a** (357.3 mg, 29%) as a colourless oil. $R_f$ = 0.46 (cyclohexane/EtOAc, 85:15). $^1$H NMR ($CDCl_3$, 360 MHz) δ (ppm): 3.70 (s, 3H, $OCH_3$), 3.65 (m, 1H, $H_c$), 2.48-2.39 (m, 2H, OH and $H_b$), 1.69 (m, 1H, $H_{a'}$), 1.66 (m, 1H, $H_{a'}$), 1.50-1.20 (m, 14H, $H_d$-$H_g$, $H_{b'}$-$H_{d'}$), 0.88 (t, 3H, *J* = 6.6 Hz, $H_h$ or $H_{e'}$), 0.87 (t, 3H, *J* = 6.5 Hz, $H_{e'}$ or $H_h$). $^{13}$C NMR ($CDCl_3$, 90 MHz) δ (ppm): 176.4 ($C_a$), 72.4 ($C_c$), 51.7 ($OCH_3$), 51.1 ($C_b$), 35.8, 31.9, 31.8, 29.7, 27.2, 25.5, 22.7, 22.6 (8C, $C_d$-$C_g$, $C_{a'}$-$C_{d'}$), 14.2, 14.1 (2C, $C_h$, $C_{e'}$). HRMS (ESI): calcd for $C_{14}H_{28}NaO_3$ $[M+Na]^+$ 267.1931, found 267.1922. $[\alpha]_D^{20}$ = + 12 (*c* 0.77, $CHCl_3$).

##### (2*R*, 3*R*)-2-pentyl-3-triethylsilyloxy-octanoic acid 14a

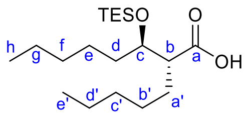


A 10% aqueous NaOH solution (2.9 mL) was added to compound **13a** (95 mg, 0.39 mmol) in MeOH (2.9 mL). The reaction mixture was stirred overnight at 45 °C and then acidified to pH ≈ 1 with an aqueous HCl solution (1M). The mixture was extracted three times with EtOAc, the combined organic layers were dried over $Na_2SO_4$ and concentrated under reduced pressure.

The residue was purified by flash silica gel chromatography (cyclohexane/EtOAc, 80:20) to give the corresponding carboxylic acid (65 mg, 73%) as a colourless oil. $R_f$ = 0.33 (cyclohexane/EtOAc, 1:1). $^1$H NMR ($CDCl_3$, 360 MHz) δ (ppm): 3.72 (m, 1H, $H_c$), 2.47 (td, 1H, *J* = 9.0 Hz, *J* = 5.3 Hz, $H_b$), 1.75 (m, 1H, $H_{a'}$), 1.62 (m, 1H, $H_{a'}$), 1.55-1.23 (m, 14H, $H_d$-$H_g$, $H_{b'}$-$H_{d'}$), 0.89 (t, 3H, *J* = 6.7 Hz, $H_h$ or $H_{e'}$), 0.88 (t, 6H, *J* = 6.7 Hz, $H_{e'}$ or $H_h$). $^{13}$C NMR ($CDCl_3$, 90 MHz) δ (ppm): 180.7 ($C_a$), 72.3 ($C_c$), 51.1 ($C_b$), 35.6, 31.8, 29.6, 27.1, 25.5, 22.7, 22.6 (8C, $C_d$-$C_g$, $C_{a'}$-$C_{d'}$), 14.2, 14.1 (2C, $C_h$, $C_{e'}$). HRMS (ESI): calcd for $C_{33}H_{26}NaO_3$ $[M+Na]^+$ 253.1774, found 253.1764. $[\alpha]_D^{20}$ = +15 (*c* 1.09, $CHCl_3$). To a solution of this compound (52 mg, 0.23 mmol) in dry pyridine (2 mL) was added dropwise TESCl (190 µL, 1.13 mmol, 5 equiv). The reaction mixture was stirred overnight at 60 °C and was then diluted with a saturated aqueous NaCl solution. The reaction mixture was extracted three times with EtOAc. The combined organic layers were dried over $Na_2SO_4$ and concentrated under reduced pressure. The residue was purified by flash silica gel chromatography (cyclohexane/EtOAc, 98:2) to give compound **14a** (29.6 mg, 38%) as a colourless oil. $R_f$ = 0.63 (cyclohexane/EtOAc 85:15). $^1$H NMR ($CDCl_3$, 360 MHz) δ (ppm): 3.87 (m, 1H, $H_c$), 2.51 (m, 1H, $H_b$), 1.71-1.59 (m, 1H, $H_{a'}$), 1.58-1.47 (m, 3H, $H_d$, $H_{a'}$), 1.40-1.22 (m, 12H, $H_e$-$H_g$, $H_{b'}$-$H_{d'}$), 0.97 (t, 9H, *J* = 7.9 Hz, $Si(CH_2C\mathit{H}_3)_3$), 0.88 (t, 6H, *J* = 6.9 Hz, $H_h$, $H_{e'}$), 0.64 (q, 6H, *J* = 7.9 Hz, $Si(C\mathit{H}_2CH_3)_3$). $^{13}$C NMR ($CDCl_3$, 90 MHz) δ (ppm): 177.6 ($C_a$), 73.8 ($C_c$), 51.0 ($C_b$), 35.4, 32.0, 31.8, 29.2, 27.4, 24.7, 22.7, 22.6 (8C, $C_d$-$C_g$, $C_{a'}$-$C_{d'}$), 14.1 (2C, $C_h$, $C_{e'}$), 6.9 ($Si(CH_2\mathit{C}H_3)_3$), 5.1 ($Si(\mathit{C}H_2CH_3)_3$). HRMS (ESI): calcd for $C_{19}H_{41}O_3Si$ $[M+H]^+$ 345.2819, found 345.2802, calcd for $C_{21}H_{40}NaO_3Si$ $[M+Na]^+$ 367.2639, found 367.2621. $[\alpha]_D^{20}$ = + 7 (*c* 1.5, $CHCl_3$).

**Methyl (2*R*, 3*R*)-2-decyl-3-hydroxy-tridecanoate 13b**

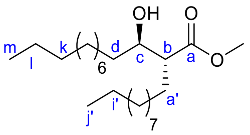


To a solution of dry diisopropylamine (722 µL, 7.16 mmol, 3.5 equiv) in THF (5.9 mL) cooled to -40 °C were added dropwise 3.84 mL of *n*-butyllithium (6.14 mmol, 1.6 M in hexane, 3 equiv). After stirring for 40 min at -40 °C, the mixture was cooled to -78 °C and a solution of β-hydroxyester **10c** (500 mg, 2.05 mmol) in dry THF (14.9 mL) was added *via* cannula. The reaction mixture was stirred for 1h30 at -78 °C then 1-iodododecane (1.31 mL, 6.14 mmol, 3 equiv) and HMPA (427 µL, 2.46 mmol, 1.2 equiv) were added slowly. The mixture was warmed slowly to -10 °C and stirred overnight at this temperature. The mixture was diluted with a saturated aqueous $NH_4Cl$ solution and extracted three times with EtOAc. The combined organic layers were washed with an aqueous NaCl solution, dried over $Na_2SO_4$ and concentrated under reduced pressure. The residue was purified by flash silica gel chromatography (cyclohexane/EtOAc, 98:2 to 95:5) to give compound **13b** (185 mg, 27%) as a colourless oil. $R_f$ = 0.47 (cyclohexane/EtOAc, 85:15). $^1$H NMR ($CDCl_3$, 300 MHz) δ (ppm): 3.70 (s, 3H, $OCH_3$), 3.65 (m, 1H, $H_c$), 2.43 (m, 1H, $H_b$), 2.35 (bs, 1H, OH), 1.79-1.51 (m, 2H, $H_{a'}$), 1.51-1.15 (m, 34H, $H_d$-$H_l$, $H_{b'}$-$H_{i'}$), 0.87 (t, 6H, *J* = 6.7 Hz, $H_m$, $H_{j'}$). $^{13}$C NMR ($CDCl_3$, 75 MHz) δ (ppm): 176.4 ($C_a$), 72.4 ($C_c$), 51.7 ($OCH_3$), 51.1 ($C_b$), 35.8, 32.0, 29.8, 29.7, 29.6, 29.5, 29.4, 27.6, 25.6, 22.8 (18C, $C_d$-$C_l$, $C_{a'}$-$C_{i'}$), 14.3 (2C, $C_m$, $C_{j'}$). HRMS (ESI): calcd for $C_{24}H_{48}NaO_3$ $[M+Na]^+$ 407.3496, found 407.3478. $[\alpha]_D^{20}$ = + 9 (*c* 1.02, $CHCl_3$).

**(2*R*, 3*R*)-2-Decyl-3-triethylsilyloxy-tridecanoic acid 14b**

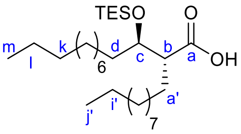


A 10% aqueous NaOH solution (6 mL) was added to compound **13b** (185 mg, 0.48 mmol) in MeOH (6 mL). The reaction mixture was stirred overnight at 45 °C and then acidified to pH ≈ 1 with an aqueous HCl solution (1M). The mixture was extracted three times with EtOAc, the combined organic layers were dried over $Na_2SO_4$ and concentrated under reduced pressure. The residue was purified by flash silica gel chromatography (cyclohexane/EtOAc, 98:2 to 95:5) to give the corresponding carboxylic acid (143.9 mg, 80%) as a colourless oil. $R_f$ = 0.05 (cyclohexane/EtOAc, 8:2). $^1$H NMR ($CDCl_3$, 300 MHz) δ (ppm): 3.71 (m, 1H, $H_c$), 2.45 (td, 1H, *J* = 8.8 Hz, *J* = 5.4 Hz, $H_b$), 1.79-1.19 (m, 36H, $H_d$-$H_l$, $H_{a'}$-$H_{i'}$), 0.88 (t, 6H, *J* = 6.8 Hz, $H_m$, $H_{j'}$). $^{13}$C NMR ($CDCl_3$, 75 MHz) δ (ppm): 180.3 ($C_a$), 72.3 ($C_c$), 51.1 ($C_b$), 35.6, 32.1, 29.8, 29.7, 29.6, 29.5, 27.4, 25.9, 22.8 (18C, $C_d$-$C_l$, $C_{a'}$-$C_{i'}$), 14.3 (2C, $C_m$, $C_{j'}$). HRMS (ESI): calcd for $C_{23}H_{46}NaO_3$ $[M+Na]^+$ 393.3339, found 393.3323. $[\alpha]_D^{20}$ = +12 (*c* 0.99, $CHCl_3$). To a solution of this compound (100 mg, 0.27 mmol) in dry pyridine (2.4 mL) was added dropwise TESCl (226 µL, 1.35 mmol, 5 equiv). The reaction mixture was stirred overnight at 60 °C and was then diluted with a saturated aqueous NaCl solution. The reaction mixture was extracted three times with EtOAc. The combined organic layers were dried over $Na_2SO_4$ and concentrated under reduced pressure. The residue was purified by flash silica gel chromatography (cyclohexane/EtOAc, 99:1 to 98:2) to give compound **14b** (59.4 mg, 45%) as a colourless oil. $R_f$ = 0.77 (cyclohexane/EtOAc 8:2). $^1$H NMR ($CDCl_3$, 300 MHz), δ (ppm): 3.84 (m, 1H, $H_c$), 2.50 (ddd, *J* = 9.2 Hz, *J* = 5.7 Hz, *J* = 2.8 Hz, 1H, $H_b$), 1.79-1.15 (m, 36H, $H_d$-$H_l$, $H_{a'}$-$H_{i'}$), 0.99 (t, 9H, *J* = 8.0 Hz, $Si(CH_2CH_3)_3$), 0.88 (t, 6H, *J* = 6.8 Hz, $H_m$, $H_{j'}$), 0.67 (q, 6H, *J* = 8.0 Hz, $Si(CH_2CH_3)_3$). $^{13}$C NMR ($CDCl_3$, 75 MHz) δ (ppm): 177.0 ($C_a$), 73.9 ($C_c$), 50.8 ($C_b$), 35.6, 32.1, 29.7, 29.6, 29.5, 29.4, 27.7, 25.1, 22.8 (18C, $C_d$-$C_l$, $C_{a'}$-$C_{i'}$), 14.3 (2C, $C_m$, $C_{j'}$), 6.9 ($Si(CH_2CH_3)_3$), 5.1 ($Si(CH_2CH_3)_3$). HRMS (ESI): calcd for $C_{29}H_{60}NaO_3Si$ $[M+Na]^+$ 507.4204, found 507.4180. $[\alpha]_D^{20}$ = + 6 (*c* 1.02, $CHCl_3$).

**(*R*)-3-Methoxy-octadecanoic acid 12**

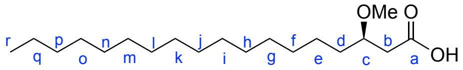


Proton sponge® (272 mg, 1.27 mmol, 4 equiv) and $Me_3O{\cdot}BF_4$ (188 mg, 1.27 mmol, 4 equiv) were added to a solution of **10b**[iii] (100 mg, 0.32 mmol) in dry $CH_2Cl_2$ (1.5 mL). The reaction mixture was stirred for 24 h at room temperature then quenched by an aqueous solution of HCl 1N to acidify the solution to pH≈1. The solution was then filtered and washed by small portions of $CH_2Cl_2$. The filtrate was diluted with $CH_2Cl_2$ and washed twice by an aqueous solution of HCl 1N and by a saturated aqueous solution of NaCl. The organic layer was dried over $Na_2SO_4$, filtered and concentrated under reduced pressure. The residue was purified by flash silica gel chromatography (cyclohexane/EtOAc 98:2) to give the expected *O*-methylated derivative (86 mg, 83%).$^1$H NMR ($CDCl_3$, 300 MHz), δ (ppm): 3.68 (s, 3H, -$OCH_3$), 3.62 (m, 1H, $H_c$), 3.33 (s, 3H, C(=O)-$OCH_3$), 2.53 (dd, 1H, *J* = 15.2 Hz, *J* = 7.3 Hz, $H_b$), 2.40 (dd, 1H, *J* = 15.2 Hz, *J* = 5.4 Hz, $H_{b'}$), 1.61-1.13 (m, 28H, $H_d$-$H_q$), 0.87 (t, 3H, *J* = 6.8 Hz, $H_r$); $^{13}$C NMR ($CDCl_3$, 75 MHz) ; δ (ppm): 172.6 ($C_a$), 78.0 ($C_c$), 57.2 (C(=O)-O$\underline{C}H_3$), 51.9 (-$OCH_3$), 39.6 ($C_b$), 39.6, 34.1, 32.2, 29.9, 29.6, 25.4, 23.0 (14C, $C_d$-$C_q$), 14.4 ($C_r$); ESI HRMS: for $C_{20}H_{40}NaO_3$ $[M+Na]^+$: calcd 351.2870, found 351,2861. A 10% aqueous NaOH solution (3 mL) was added to a solution of this compound (95 mg, 0.29 mmol) in MeOH (3 mL). The reaction mixture was then stirred for 4 h at 40°C. Aqueous solution of HCl (6 M) was added to acidify the solution to pH ≈ 1. The solution was extracted three times with EtOAc and the combined organic layers were dried over $Na_2SO_4$, filtered and concentrated under reduced pressure. The desired product **12** (75 mg, 82%) was obtained as a colorless oil. $^1$H NMR ($CDCl_3$, 300 MHz); δ (ppm): 3.67 (m, 1H, $H_c$), 3.43 (s, 3H, -$OCH_3$), 2.60 (dd, 1H, *J* = 15.6 Hz, *J* = 6.7 Hz, $H_b$), 2.54 (dd, 1H, *J* = 15.6 Hz, *J* = 5.3 Hz, $H_{b'}$), 1.74-1.16 (m, 28H, $H_d$-$H_q$), 0.92 (t, 3H, *J* = 6.8 Hz, $H_r$); $^{13}$C NMR ($CDCl_3$, 75 MHz); δ (ppm): 176.3 ($C_a$), 77.8 ($C_c$), 57.1 (C(=O)-O<u>C</u>$H_3$), 39.2 ($C_b$), 39.2, 33.7, 32.2, 29.9, 29.8, 29.6, 25.2, 22.9 (14C, $C_d$-$C_q$), 14.36 ($C_r$); ESI HRMS: for $C_{19}H_{38}NaO_3$ $[M+Na]^+$: calcd 337.2713, found 337.2701.

**6-*O*-((2*R*, 3*R*)-2-Decyl-3-hydroxy-tridecanoyl)-α,α-D-trehalose 1**

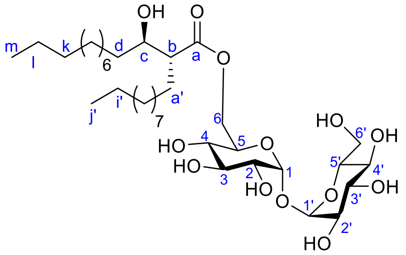

A solution of compound **15**[x] (183 mg, 0.24 mmol, 2 equiv) in DCM (3 mL) was added to compound **14b** (57.3 mg, 0.12 mmol). EDCI (45.3 mg, 0.24 mmol, 2 equiv) and DMAP (28.9 mg, 0.24 mmol, 2 equiv) were then added and the reaction mixture was stirred at room temperature overnight. Saturated aqueous NaCl solution was added, and the reaction mixture was extracted three times with DCM. The combined organic layers were dried over $Na_2SO_4$ and concentrated under reduced pressure. The residue was purified by flash silica gel chromatography (cyclohexane/EtOAc, 99:1 to 90:10) to give the expected mono-*O*-esterified product (45.9 mg, 31%) as a colourless oil. $R_f$ = 0.58 (cyclohexane/EtOAc 9:1). $^1$H NMR ($CDCl_3$, 300 MHz) δ (ppm): 4.90 (d, 1H, *J* = 3.1 Hz, $H_1$ or $H_{1'}$), 4.84 (d, 1H, *J* = 3.1 Hz, $H_{1'}$ or $H_1$), 4.35 (dd, 1H, *J* = 11.7 Hz, *J* = 2.1 Hz, $H_{6a}$), 4.05 (dd, 1H, *J* = 11.7 Hz, *J* = 3.7 Hz $H_{6b}$), 4.01-3.80 (m, 5H, $H_c$, $H_3$, $H_{3'}$, $H_5$, $H_{5'}$), 3.71-3.66 (m, 2H, $H_{6a'}$, $H_{6b'}$), 3.50 (at, 1H, *J* = 8.9 Hz, $H_4$ or $H_{4'}$), 3.47 (at, 1H, *J* = 8.8 Hz, $H_{4'}$ or $H_4$), 3.42 (dd, 1H, *J* = 9.3 Hz, *J* = 3.1 Hz, $H_2$ or $H_{2'}$), 3.38 (dd, 1H, *J* = 9.3 Hz, *J* = 3.1 Hz, $H_{2'}$ or $H_2$), 2.53 (ddd, *J* = 10.6 Hz, *J* = 5.5 Hz, *J* = 3.2 Hz, 1H, $H_b$), 1.80-1.09 (m, 36H, $H_d$-$H_l$, $H_{a'}$-$H_{i'}$), 0.96 (t, 9H, *J* = 7.8 Hz, Si($CH_2C\mathit{H}_3$)$_3$), 0.88 (t, 3H, *J* = 6.6 Hz, $H_m$ or $H_{j'}$), 0.87 (t, 3H, *J* = 6.5 Hz, $H_{j'}$ or $H_m$), 0.60 (q, 6H, *J* = 7.9 Hz, Si($C\mathit{H}_2CH_3$)$_3$), 0.17-0.10 (m, 54H, 6 Si($CH_3$)$_3$). $^{13}$C NMR ($CDCl_3$, 75 MHz) δ (ppm): 174.2 ($C_a$), 94.7, 94.6 (2C, $C_1$, $C_{1'}$), 73.6, 73.5, 73.4, 73.0, 72.9, 72.8 (6C, $C_3$, $C_{3'}$, $C_5$, $C_{5'}$, $C_2$, $C_{2'}$), 72.0, 71.5 (2C, $C_4$, $C_{4'}$), 70.8 ($C_c$), 62.6 ($C_6$), 61.8 ($C_{6'}$), 52.5 ($C_b$), 33.6 ($C_d$), 32.1, 30.0, 29.9, 29.8, 29.7, 29.6, 29.5, 29.4, 28.3, 26.2, 25.4, 22.8 (17C, $C_e$-$C_l$, $C_{a'}$-$C_{i'}$), 14.3 (2C, $C_m$, $C_{j'}$), 7.1 (Si($CH_2\mathit{C}H_3$)$_3$), 5.2 (Si($\mathit{C}H_2CH_3$)$_3$), 1.2, 1.1, 1.0, 0.9, 0.3, 0.2 (6 Si($CH_3$)$_3$). HRMS (ESI): calcd for $C_{59}H_{128}NaO_{13}Si_7$ $[M+Na]^+$ 1263.7632, found 1263.7625. $[\alpha]_D^{20}$ = + 61 (*c* 0.83, $CHCl_3$). To a solution of this compound (45.9 mg, 0.037 mmol) in MeOH (3.4 mL) were added 350 mg of Dowex 50WX8 ($H^+$ form). The reaction mixture was stirred at room temperature for 1h15, then filtered, washed three times with MeOH and concentrated under reduced pressure. The residue was purified by flash silica gel chromatography (EtOAc/MeOH, 95:5 to 90:10) to give the desired TMM analog **1** (20 mg, 78%) as a colourless oil. $R_f$ = 0.71 (EtOAc/MeOH, 10 mL, 7:3 + 2 drops of water). $^1$H NMR ($CD_3OD$, 300 MHz) δ (ppm): 5.09 (d, 2H, *J* = 3.7 Hz, $H_1$ and $H_{1'}$), 4.62 (s, 1H, OH), 4.47 (dd, 1H, *J* = 11.8 Hz, *J* = 1.7 Hz, $H_{6a}$), 4.18 (dd, 1H, *J* = 11.8 Hz, *J* = 5.3 Hz $H_{6b}$), 4.07 (ddd, 1H, *J* = 9.9 Hz, *J* = 5.3 Hz, *J* = 1.7 Hz, $H_5$) 3.87-3.75 (m, 4H, $H_{5'}$, $H_{6a'}$, $H_3$, $H_{3'}$), 3.73-3.64 (m, 2H, $H_{6b'}$, $H_c$), 3.48 (dd, 2H, *J* = 9.7 Hz, *J* = 3.7 Hz, $H_2$ and $H_{2'}$), 3.39-3.32 (m, 2H, $H_4$, $H_{4'}$), 2.43 (ddd, 1H, *J* = 10.2 Hz, *J* = 7.3 Hz, *J* = 4.1 Hz, $H_b$), 1.70-1.17

(m, 36H, $H_d$-$H_l$, $H_{a'}$-$H_{i'}$), 0.90 (t, 6H, $J$ = 6.9 Hz, $H_m$ and $H_{j'}$). $^{13}C$ NMR ($CD_3OD$, 75 MHz) δ (ppm): 176.2 ($C_a$), 95.3, 95.2 (2C, $C_1$, $C_{1'}$), 74.4, 74.3, 73.8 (3C, $C_3$, $C_{3'}$, $C_{5'}$), 73.6 ($C_c$), 73.2 (2C, $C_2$, $C_{2'}$), 72.0, 71.9 (2C, $C_4$, $C_{4'}$), 71.4 ($C_5$), 64.4 ($C_6$), 62.6 ($C_{6'}$), 54.2 ($C_b$), 35.6, 33.1, 30.8, 30.7, 30.6, 30.5, 29.8, 29.7, 28.6, 26.6, 23.8 (18C, $C_d$-$C_l$, $C_{a'}$-$C_{i'}$), 14.5 (2C, $C_m$, $C_{j'}$). HRMS (ESI): calcd for $C_{35}H_{66}NaO_{13}$ $[M+Na]^+$ 717.4396, found 717.4365. $[\alpha]_D^{20}$ = + 93 (*c* 1, $CHCl_3$).

**6-*O*-((2*R*, 3*R*)-2-Pentyl-3-hydroxy-octanoyl)-α,α-D-trehalose 2**

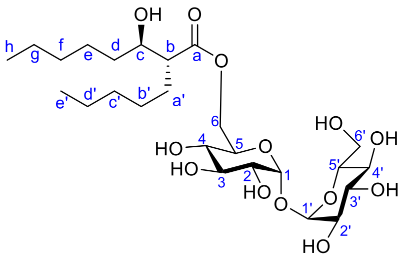


A solution of compound **14a** (46 mg, 0.13 mmol) in DCM (0.5 mL) was added to a solution of dried EDCI (32.6 mg, 0.26 mmol, 2 equiv) and DMAP (12.8 mg, 0.07 mmol, 0.5 equiv) in DCM (0.4 mL). A solution of compound **15**[x] (103.3 mg, 0.13 mmol, 1 equiv) in DCM (1.2 mL) was added and the reaction mixture was stirred at room temperature for 4 h. Then 2 more equiv of EDCI and 0.5 equiv of DMAP were added and the solution was stirred at RT for 2 h. Finally, DCC (110.2 mg, 4 equiv) and DMAP (12.8 mg, 0.5 equiv) were added, and the reaction mixture was stirred at room temperature overnight then filtered, washed three times with cold DCM and concentrated under reduced pressure. The residue was purified by flash silica gel chromatography (cyclohexane/EtOAc, 99:1 to 98:2) to give the mono-*O*-esterified product (81.7 mg, 56%) as a colourless oil. $R_f$ = 0.55 (cyclohexane/EtOAc 9:1). $^1H$ NMR ($CDCl_3$, 360 MHz) δ (ppm): 4.90 (d, 1H, $J$ = 3.2 Hz, $H_1$ or $H_{1'}$), 4.84 (d, 1H, $J$ = 3.2 Hz, $H_{1'}$ or $H_1$), 4.35 (dd, 1H, $J$ = 11.9 Hz, $J$ = 2.2 Hz, $H_{6a}$), 4.06 (dd, 1H, $J$ = 11.9 Hz, $J$ = 3.6 Hz $H_{6b}$), 4.01-3.94 (m, 2H, $H_c$, $H_5$), 3.90 (at, 1H, $J$ = 9.0 Hz, $H_3$ or $H_{3'}$), 3.89 (t, 1H, $J$ = 9.0 Hz, $H_{3'}$ or $H_3$), 3.83 (td, 1H, $J$ = 9.4 Hz, $J$ = 3.5 Hz, $H_{5'}$), 3.71-3.66 (m, 2H, $H_{6a'}$, $H_{6b'}$), 3.50 (at, 1H, $J$ = 9.0 Hz, $H_4$ or $H_{4'}$), 3.46 (at, 1H, $J$ = 9.0 Hz, $H_{4'}$ or $H_4$), 3.42 (dd, 1H, $J$ = 9.0 Hz, $J$ = 3.2 Hz, $H_2$ or $H_{2'}$), 3.38 (dd, 1H, $J$ = 9.0 Hz, $J$ = 3.0 Hz, $H_{2'}$ or $H_2$), 2.54 (ddd, $J$ = 10.8 Hz, $J$ = 5.4 Hz, $J$ = 3.2 Hz, 1H, $H_b$), 1.75 (dd, 1H, $J$ = 7.4 Hz, $J$ = 5.4 Hz, OH), 1.65-1.16 (m, 16H, $H_d$-$H_g$, $H_{a'}$-$H_{d'}$), 0.96 (t, 9H, $J$ = 7.9 Hz, $Si(CH_2CH_3)_3$), 0.88 (t, 3H, $J$ = 7.0 Hz, $H_h$ or $H_{e'}$), 0.87 (t, 3H, $J$ = 6.7 Hz, $H_{e'}$ or $H_h$), 0.60 (q, 6H, $J$ = 7.9 Hz, $Si(CH_2CH_3)_3$), 0.17-0.10 (m, 54H, 6 $Si(CH_3)_3$). $^{13}C$ NMR ($CDCl_3$, 90 MHz) δ (ppm): 174.1 ($C_a$), 94.6, 94.5 (2C, $C_1$, $C_{1'}$), 73.6, 73.5, 73.4 (3C, $C_3$, $C_{3'}$, $C_5$), 73.0, 72.9, 72.8 (3C, $C_2$, $C_{2'}$, $C_{5'}$), 72.0, 71.5 (2C, $C_4$, $C_{4'}$), 70.8 ($C_c$), 62.6 ($C_6$), 61.8 ($C_{6'}$), 52.5 ($C_b$), 35.6, 32.1, 32.0, 28.0, 26.1, 25.0, 22.8, 22.7 (8C, $C_d$-$C_g$, $C_{a'}$-$C_{d'}$), 14.2 (2C, $C_h$, $C_{e'}$), 7.1 ($Si(CH_2CH_3)_3$), 5.2 ($Si(CH_2CH_3)_3$), 1.2, 1.1, 1.0, 0.3, 0.2 (6 $Si(CH_3)_3$). HRMS (ESI): calcd for $C_{49}H_{108}NaO_{13}Si_7$ $[M+Na]^+$ 1123.6067, found 1123.6012. $[\alpha]_D^{20}$ = + 59 (*c* 0.9, $CHCl_3$). To a solution of this compound (81 mg, 0.07 mmol) in MeOH (4 mL) were added 200 mg of Dowex 50WX8 ($H^+$ form). The reaction mixture was stirred at room temperature for 1h15, then filtered, washed three times with MeOH and concentrated under reduced pressure. The residue was purified by flash silica gel chromatography (EtOAc/MeOH, 90:10) to give the desired product **2** (35 mg, 86%) as a colourless oil. $R_f$ = 0.32 (EtOAc/MeOH, 10 mL, 7:3 + 2 drops of water). $^1H$ NMR ($CD_3OD$, 360 MHz) δ (ppm): 5.05 (d, 2H, $J$ = 3.6 Hz, $H_1$ and $H_{1'}$), 4.44 (dd, 1H, $J$ = 11.8 Hz, $J$ = 1.9 Hz, $H_{6a}$), 4.14 (dd, 1H, $J$ = 11.8 Hz, $J$ = 5.0 Hz, $H_{6b}$), 4.03 (ddd, 1H,

$J$ = 10.1 Hz, $J$ = 5.0 Hz, $J$ = 1.9 Hz, $H_5$) 3.83-3.74 (m, 4H, $H_3$, $H_{3'}$, $H_{5'}$, $H_{6a'}$), 3.68-3.62 (m, 2H, $H_{6b'}$, $H_c$), 3.45 (dd, 2H, $J$ = 9.6 Hz, $J$ = 3.5 Hz, $H_2$, $H_{2'}$), 3.36-3.26 (m, 2H, $H_4$, $H_{4'}$), 2.41 (ddd, 1H, $J$ = 10.7 Hz, $J$ = 6.8 Hz, $J$ = 4.7 Hz, $H_b$), 1.65-1.19 (m, 16H, $H_d$-$H_g$, $H_{a'}$-$H_{d'}$), 0.89 (t, 3H, $J$ = 6.9 Hz, $H_h$ or $H_{e'}$), 0.87 (t, 3H, $J$ = 6.9 Hz, $H_{e'}$ or $H_h$). $^{13}C$ NMR ($CD_3OD$, 90 MHz) δ (ppm): 176.1 ($C_a$), 95.3, 95.2 (2C, $C_1$, $C_{1'}$), 74.4, 74.3, 73.8 (3C, $C_3$, $C_{3'}$, $C_5$), 73.6 ($C_c$), 73.2, 73.1 (2C, $C_2$, $C_{2'}$), 72.0, 71.9 (2C, $C_4$, $C_{4'}$), 71.4 ($C_{5'}$), 64.3 ($C_6$), 62.6 ($C_{6'}$), 54.3 ($C_b$), 35.6, 33.0, 32.9, 29.8, 28.3, 26.4, 23.7, 23.5 (8C, $C_d$-$C_g$, $C_{a'}$-$C_{d'}$), 14.5, 14.4 (2C, $C_h$, $C_{e'}$). HRMS (ESI): calcd for $C_{25}H_{46}NaO_{13}$ $[M+Na]^+$ 577.2831, found 577.2813. $[\alpha]_D^{20}$ = + 111 ($c$ 0.86, MeOH/$CHCl_3$ 3/1).

##### 6-*O*-(2-Tetradecyl-octadecanoyl)-α,α-D-trehalose 3

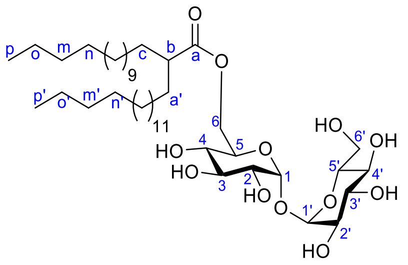


To a solution of compound **15**[x] (242 mg, 0.31 mmol, 2 equiv) in DCM (1 mL) were added, 64 mg of DCC (0.45 mmol, 2 equiv) and 30 mg of DMAP (0.16 mmol, 1 equiv) and 75 mg of compound **8** (0.156 mmol, 1 equiv) diluted with 2 mL of DCM. The reaction mixture was stirred at room temperature for 6 h. Then, the mixture was diluted with DCM and a saturated aqueous NaCl solution was added. The resulted mixture was extracted three times with DCM. The combined organic layers were dried over $Na_2SO_4$ and concentrated under reduced pressure. The residue was filtered using flash silica gel chromatography (cyclohexane to cyclohexane: EtOAc, 13:1) to give the esterified intermediate (41.4 mg, 21%). $^1H$ NMR ($CDCl_3$, 300 MHz) δ (ppm): 4.92 (d, 1H, $J$ = 3.2 Hz, $H_1$ or $H_{1'}$), 4.85 (d, 1H, $J$ = 3.0 Hz, $H_{1'}$ or $H_1$), 4.50 (dd, 1H, $J$ = 12.8 Hz, $J$ = 3.1 Hz, $H_{6a}$), 4.03-3.95 (m, 2H, $H_5$, $H_{6b}$), 3.90 (at, 1H, $J$ = 9.0 Hz, $H_3$), 3.89 (at, 1H, J = 9.0 Hz, $H_{3'}$), 3.83 (m, 1H, $H_{5'}$), 3.75-3.61 (m, 2H, $H_{6a'}$, $H_{6b'}$), 3.49 (at, 1H, $J$ = 9.0 Hz, $H_4$), 3.47 (at, 1H, $J$ = 9.0 Hz, $H_{4'}$), 3.42 (dd, 1H, $J$ = 9.0 Hz, $J$ = 3.2 Hz, $H_2$), 3.37 (dd, 1H, $J$ = 9.0 Hz, $J$ = 3.0 Hz, $H_{2'}$), 2.35 (m, 1H, $H_b$), 1.90-0.95 (m, 56H, 28 $CH_2$), 0.88 (2 t, 6H, $J$ = 6.9 Hz, 2 $CH_3$), 0.20-0.07 (m, 54H, 6 $Si(CH_3)_3$); $^{13}C$ NMR ($CDCl_3$, 75 MHz) δ (ppm): 176.5 (CO), 94.6 ($C_1$), 94.5 ($C_{1'}$), 73.7, 73.5, 73.1, 73.0, 72.8, 72.1, 71.5, 71.1 ($C_2$, $C_{2'}$, $C_3$, $C_{3'}$, $C_4$, $C_{4'}$, $C_5$, $C_{5'}$), 62.2 ($C_6$), 61.8 ($C_{6'}$), 46.0 ($C_b$), 32.1, 29.9, 29.8, 29.7, 29.5, 29.4, 27.6, 27.5 (28C, $C_c$-$C_o$, $C_{a'}$-$C_{o'}$), 14.3 (2C, $C_p$, $C_{p'}$), 1.2, 1.1, 1.0, 0.3, 0.2 (6 $Si(CH_3)_3$). HRMS (ESI): calcd for $C_{62}H_{132}NaO_{12}$ $[M+Na]^+$ 1259.8227, found 1259.8184. The filtered derivative was therefore diluted with MeOH (5 mL), and 100 mg of Dowex 50WX8 ($H^+$ form) was added to the solution. The reaction mixture was stirred at room temperature for 40 min, then filtered, washed three times with MeOH and concentrated under reduced pressure. The residue was purified by flash silica gel chromatography (DCM/MeOH, 85:15) to give the desired product **3** (21.1 mg, 77%) as a white solid. $R_f$ = 0.26 (DCM/MeOH, 81:15). $^1H$ NMR ($CD_3OD$/$CDCl_3$, 360 MHz) δ (ppm): 5.10 (m, 2H, $H_1$, $H_{1'}$), 4.39 (dd, 1H, $J$ = 11.9 Hz, $J$ = 2.2 Hz, $H_{6a}$), 4.22 (dd, 1H, $J$ = 11.9 Hz, $J$ = 4.3 Hz, $H_{6b}$), 3.99 (ddd, 1H, $J$ = 10.0 Hz, $J$ = 4.3 Hz, $J$ = 2.2 Hz, $H_5$), 3.84-3.73 (m, 4H, $H_3$, $H_{3'}$, $H_{5'}$, $H_{6a'}$), 3.64 (dd, 1H, $J$ = 12.2 Hz, $J$ = 5.7 Hz, $H_{6b'}$), 3.49 (dd, 1H, $J$ = 9.7 Hz, $J$ = 3.2 Hz, $H_2$ or $H_{2'}$), 3.48 (dd, 1H $J$ = 10.0 Hz, $J$ = 3.6 Hz, $H_{2'}$ or $H_2$), 3.42-3.30 (m, 2H, $H_4$, $H_{4'}$), 2.35 (m, 1H, $H_b$), 1.65-1.1 (m, 56H, $H_c$-$H_o$ and $H_{a'}$-$H_{o'}$), 0.89 (2t, 6H, $J$ = 6.8 Hz, $H_p$ and $H_{p'}$).

$^{13}$C NMR ($CD_3OD/CDCl_3$, 75 MHz) δ (ppm): 178.1 ($C_a$), 94.7, 94.6 (2C, $C_1$, $C_{1'}$), 74.3, 74.0, 73.3, 72.7, 72.7, 71.6; 71.4, 71.0 (8C, $C_2$, $C_{2'}$, $C_3$, $C_{3'}$, $C_4$, $C_{4'}$, $C_5$, $C_{5'}$), 63.7 ($C_6$), 62.5 ($C_{6'}$), 46.6 ($C_b$), 33.0, 32.7, 30.5, 30.4, 30.3, 30.2, 28.2, 28.1, 23.5 (28C, $C_c$-$C_o$ and $C_{a'}$-$C_{o'}$), 14.5 (2C, $C_p$, $C_{p'}$). HRMS (ESI): calcd for $C_{44}H_{84}NaO_{12}$ $[M+Na]^+$ 827.5963, found 827.5854.

##### 6-*O*-((*R*)-3-Hydroxy-octadecanoyl)-α,α-D-trehalose 4

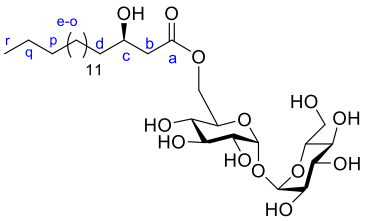


A solution of compound **15**$^x$ (132 mg, 0.17 mmol, 1.2 equiv) in $CH_2Cl_2$ (1.5 mL) was added *via* cannula to a solution of **11** (58 mg, 0.14 mmol), EDCI (43 mg, 0.28 mmol, 2.0 equiv), and DMAP (34 mg, 0.28 mmol, 2.0 equiv) in $CH_2Cl_2$ (0.5 mL). The reaction mixture was stirred overnight at room temperature and then diluted by a saturated aqueous solution of NaCl. The solution was extracted three times by $CH_2Cl_2$ and the combined organic layers were dried over $Na_2SO_4$, filtered and concentrated under reduced pressure. The residue was purified by flash silica gel chromatography (cyclohexane/EtOAc 98:2 to 96:4) to give the esterified compound (80 mg, 49%) as a colorless oil. $^1$H NMR ($CDCl_3$, 300 MHz); δ (ppm): 4.94 (d, 1H, $J$ = 3.0 Hz, $H_1$), 4.91 (d, 1H, $J$ = 3.0 Hz, $H_{1'}$), 4.30 (dd, 1H, $J$ = 11.5 Hz, $J$ = 1.7 Hz, $H_{6a}$), 4.17 (m, 1H, $H_c$), 4.09 (dd, 1H, $J$ = 11.5 Hz, $J$ = 4.8 Hz, $H_{6b}$), 4.02 (m, 1H, $H_5$), 3.97-3.88 (m, 2H, $H_3$ and $H_3$'), 3.86 (m, 1H, $H_5$'), 3.78-3.64 (m, 2H, $H_{6a}$' and $H_{6b}$'), 3.54-3.41 (m, 4H, $H_2$, $H_2$', $H_4$, $H_4$'), 2.55 (dd, 1H, $J$ = 7.0 Hz, $J$ = 15.4 Hz, 1H of $H_b$), 2.46 (dd, 1H, $J$ = 6.2 Hz, $J$ = 15.4 Hz, 1H of $H_b$), 1.75 (at, 1H, $J$ = 6.6 Hz, OH), 1.49 (m, 2H, m, $H_d$), 1.40-1.19 (m, 24H, $H_e$-$H_q$), 0.96 (t, 9H, $J$ = 7.9 Hz, $Si(CH_2C\mathit{H}_3)_3$), 0.90 (t, 3H, $J$ = 6.5 Hz, $H_r$), 0.61 (q, 6H, $J$ = 7.9 Hz, $Si(C\mathit{H}_2CH_3)_3$), 0.21-0.10 (54H, m, 6 $Si(CH_3)_3$); $^{13}$C NMR ($CDCl_3$, 75 MHz); δ (ppm): 171.8 (Ca), 94.5, 94.3 (C1, C1'), 73.4, 73.4, 72.9, 72.8, 72.6, 72.0, 71.4, (C2, C2', C3, C3', C4, C4', C5'), 70.7 (C5), 69.2 (Cc), 63.4 (C6), 61.7 (C6'), 42.7 (Cb), 37.7 (Cd), 31.9, 29.7, 29.7, 29.6, 29.6, 29.4, 25.2, 22.7 (Ce-Cq), 14.1 (Cr), 7.0 ($Si(CH_2\mathit{C}H_3)_3$), 4.9 ($Si(\mathit{C}H_2CH_3)_3$), 1.0, 1.0, 0.9, 0.9, 0.2, 0.1 ($Si(CH_3)_3$); ESI HRMS: calcd for $C_{54}H_{118}NaO_{13}Si_7[M+Na]^+$: 1193.6850, found 1193.6806. Dowex 50WX8 ($H^+$ form) resin (700 mg) was added to solution of the esterified compound (78 mg, 0.07 mmol) in MeOH (5 mL). The solution was stirred for 40 min at room temperature, then filtered, washed with MeOH and concentrated under reduced pressure. The residue was purified by flash silica gel chromatography ($CH_2Cl_2$/MeOH 95:5 to 90:10) to give compound **4** (41 mg, quant.) as a white solid. $^1$H NMR ($CD_3OD$ /$CDCl_3$, 360 MHz) δ (ppm): 5.11, 5.10 (2d, 2H, $J$ = 3.8 Hz, $H_1$ and $H_{1'}$), 4.45 (dd, 1H, $J$ = 11.9 Hz, $J$ = 2.0 Hz, $H_{6a}$), 4.22 (dd, 1H, $J$ = 11.9 Hz, $J$ = 5.4 Hz, $H_{6b}$), 4.09 (m, 1H, $H_5$), 4.02 (m, 1H, $H_c$), 3.84-3.78 (m, 4H, $H_3$, $H_{3'}$, $H_{6a'}$, $H_{5'}$), 3.70 (dd, 1H, $J$ = 12.0 Hz, $J$ = 5.7 Hz $H_{6b'}$), 3.51 (dd, 1H, $J$ = 9.7 Hz, $J$ = 4.1 Hz, $H_2$ or $H_{2'}$), 3.50 (dd, 1H, $J$ = 9.7 Hz, $J$ = 3.8 Hz, $H_{2'}$ or $H_2$), 3.39-3.35 (m, 2H, $H_4$, $H_{4'}$), 2.55 (dd, 1H, $J$ = 15.0 Hz, $J$ = 4.6 Hz, 1H of $H_b$), 2.45 (dd, 1H, $J$ = 15.0 Hz, $J$ = 8.4 Hz, 1H of $H_b$), 1.54-1.24 (m, 28H, $H_d$-$H_q$), 0.93 (t, 3H, $J$ = 6.8 Hz, $H_r$). $^{13}$C NMR ($CD_3OD$, 90 MHz) δ (ppm): 172.0 ($C_a$), 93.9, 93.8 (2C, $C_1$, $C_{1'}$), 73.1 ($C_5$), 72.5, 71.8 (2C, $C_3$, $C_{3'}$), 70.5 (2C, $C_2$, $C_2$'), 70.0 (2C, $C_4$, $C_4$'), 69.9 ($C_5$'), 68.0 ($C_c$), 63.2 ($C_6$), 61.2 ($C_{6'}$), 41.9 ($C_b$), 36.7 ($C_d$), 31.7, 29.4, 29.3, 29.1, 25.3, 22.3 (13C, $C_e$-$C_q$), 13.0 ($C_r$). HRMS (ESI): calcd for $C_{30}H_{56}NaO_{13}$ $[M+Na]^+$ 647.3613, found 647.3614. $[\alpha]_D^{20}$ = +97 (*c* 1.27, $CHCl_3$/MeOH 1/1).

**6-*O*-(*R*)-3-methoxy-octadecanoyl)- α,α-D-trehalose 5**

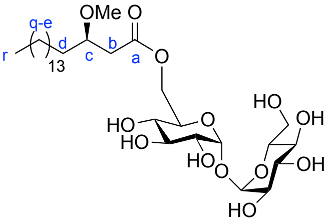

A solution of **15**$^x$ (171 mg, 0.22 mmol, 1.2 equiv) in $CH_2Cl_2$ (1.2 mL) was added *via* cannula to a solution of **12** (57 mg, 0.18 mmol), EDCI (56 mg, 0.36 mmol, 2 equiv), and DMAP (44 mg, 0.36 mmol, 2 equiv) in $CH_2Cl_2$ (0.5 mL). The reaction mixture was stirred overnight at room temperature and then diluted by a saturated aqueous solution of NaCl. The solution was extracted three times by $CH_2Cl_2$ and the combined organic layers were dried over $Na_2SO_4$, filtered and concentrated under reduced pressure. The residue was purified by flash silica gel chromatography (cyclohexane/EtOAc 98:2 to 92:8) to give the desired esterified compound (67 mg, 35%) as a colorless oil. $^1$H NMR ($CDCl_3$, 300 MHz); δ (ppm): 4.93 (d, 1H, *J* = 3.1 Hz, $H_{1'}$), 4.91 (d, 1H, *J* = 3.1 Hz, $H_1$), 4.34 (dd, 1H, *J* = 2.0 Hz, *J* = 11.7 Hz, $H_{6a}$), 4.10 (dd, 1H, *J* = 5.0 Hz, *J* = 11.7 Hz, $H_{6b}$), 4.02 (ddd, 1H, *J* = 2.0 Hz, *J* = 5.0 Hz, *J* = 9.0 Hz, $H_5$), 3.93 (at, 1H, *J* = 9.0 Hz, $H_3$), 3.91 (at, 1H, *J* = 9.0 Hz, $H_{3'}$), 3.85 (m, 1H, $H_{5'}$), 3.77-3.61 (m, 3H, $H_{6'a}$, $H_{6'b}$ and $H_c$), 3.49 (at, 1H, *J* = 9.0 Hz, $H_4$), 3.46 (at, 1H, *J* = 9.0 Hz, $H_{4'}$), 3.45 (dd, 1H, *J* = 3.1 Hz, *J* = 9.0Hz, $H_2$), 3.44 (dd, 1H, *J* = 3.1 Hz, *J* = 9.0 Hz, $H_{2'}$), 3.36 (s, 3H, -$OCH_3$), 2.63 (dd, 1H, *J* = 6.9 Hz, *J* = 15.8 Hz, 1H of $H_b$), 2.44 (dd, 1H, *J* = 5.6 Hz, *J* = 15.8 Hz, 1H of $H_b$) 1.58-1.20 (m, 28H, $H_d$-$H_q$), 0.18-0.13 (6s, 54H, 6 $Si(CH_3)_3$) ; $^{13}$C NMR ($CDCl_3$, 75 MHz); δ (ppm): 172.0 ($C_a$), 94.7 ($C_1$ or $C_{1'}$), 94.5 ($C_1$ or $C_{1'}$), 77.4 ($C_c$), 73.7, 73.5, 73.1, 73.0, 72.8, 72.2, 71.6, 70.1 (8C, $C_2$, $C_{2'}$, $C_3$, $C_{3'}$, $C_4$, $C_{4'}$, $C_5$, $C_{5'}$), 63.7 ($C_6$), 61.9 ($C_{6'}$), 57.2 (O-$CH_3$), 39.4 ($C_b$), 34.2, 32.1, 29.9, 29.6, 25.3, 22.8 (14C, $C_d$-$C_q$), 14.3 ($C_r$), 1.3, 1.2, 1.1, 1.1, 0.7, 0.3 (6 $Si(CH_3)_3$) ; ESI HRMS: calcd for $C_{49}H_{106}NaO_{13}Si_6$ $[M+Na]^+$: 1093.6141, found 1093.6101. Dowex 50WX8 ($H^+$ form) resin (480 mg) was added to a solution of the esterified compound (51 mg, 0.05 mmol) in MeOH (4 mL). The solution was stirred for 1h at room temperature, then filtered, washed with MeOH and concentrated under reduced pressure. The residue was purified by flash silica gel chromatography ($CH_2Cl_2$/MeOH 95:5 to 90:10) to give the desired product **5** (24 mg, quant.) as a white solid. $^1$H NMR ($CD_3OD$, 250 MHz); δ (ppm): 5.12 (d, 1H, *J* = 3.6 Hz, $H_1$), 5.10 (d, 1H, *J* = 3.6 Hz, $H_{1'}$), 4.41 (dd, 1H, *J* = 2.7 Hz, *J* = 11.9 Hz, $H_{6a}$), 4.23 (dd, 1H, *J* = 5.1 Hz, *J* = 11.9 Hz, $H_{6b}$), 4.07 (m, 1H, $H_5$), 3.90-3.75 (m, 4H, $H_3$, $H_{3'}$, $H_{5'}$ and $H_{6'a}$), 3.75-3.62 (m, 2H, $H_{6'b}$, $H_c$), 3.55-3.45 (m, 2H, $H_2$, $H_{2'}$), 3.37 (s, 3H, O-$CH_3$), 3.42-3.29 (m, 2H, $H_4$, $H_{4'}$), 2.58-2.50 (m, 2H, $H_b$), 1.66-1.19 (m, 28H, $H_d$-$H_q$), 0.93 (t, 3H, *J* = 6.3 Hz, $H_r$) ; $^{13}$C NMR ($CD_3OD$, 62.5 MHz); δ (ppm): 171.9 ($C_a$), 93.9, 93.8 ($C_1$, $C_{1'}$), 77.8 ($C_c$), 73.2, 73.1, 72.5, 71.8, 70.5 (7C, $C_2$, $C_{2'}$, $C_3$, $C_{3'}$, $C_4$, $C_{4'}$, $C_{5'}$), 70.0 ($C_5$), 63.2 ($C_6$), 61.3 ($C_{6'}$), 55.9 ($CH_3$-O), 38.8 ($C_b$), 33.5, 31.7, 29.4, 29.3, 29.1, 24.8, 22.3 (14C, $C_d$-$C_q$), 13.1 ($C_r$); ESI HRMS: calcd for $C_{31}H_{58}NaO_{13}$ $[M+Na]^+$: 661.3770, found 661.3751.

**6-*O*-(hexadecanoyl)-α,α-D-trehalose 6**

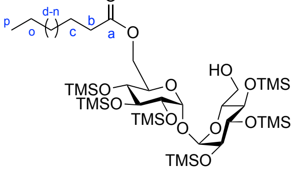

A solution of **15**$^x$ (248 mg, 0.32 mmol, 1.2 equiv) in $CH_2Cl_2$ (2 mL) was added *via* cannula to a solution of commercially available palmitic acid (70 mg, 0.27 mmol), EDCI (84 mg, 0.54

mmol, 2 equiv), and DMAP (66 mg, 0.54 mmol, 2 equiv) in $CH_2Cl_2$ (2 mL). The reaction mixture was stirred overnight at room temperature and then diluted by a saturated aqueous solution of NaCl. The solution was extracted three times by $CH_2Cl_2$ and the combined organic layers were dried over $Na_2SO_4$, filtered and concentrated under reduced pressure. The residue was purified by flash silica gel chromatography (cyclohexane/EtOAc 98:2 to 9:1) to give the esterified product (139 mg, 51%) as a colorless oil. $^1H$ NMR ($CDCl_3$, 400 MHz); δ (ppm): 4.94 (d, 1H, $J$ = 2.9 Hz, $H_1$), 4.93 (d, 1H, $J$ = 2.9 Hz, $H_{1'}$), 4.32 (dd, 1H, $J$ = 11.9 Hz, $J$ = 2.2 Hz, $H_{6a}$), 4.08 (dd, 1H, $J$ = 11.9 Hz, $J$ = 4.3 Hz, $H_{6b}$), 4.05-4.00 (m, 1H, $H_5$), 3.93 (at, 1H, $J$ = 9.0 Hz, $H_3$), 3.91 (at, 1H, $J$ = 9.0 Hz, $H_{3'}$), 3.86 (m, 1H, $H_{5'}$), 3.77-3.63 (m, 2H, $H_{6'a}$ and $H_{6'b}$), 3.50 (at, 1H, $J$ = 9.0 Hz, $H_4$), 3.49 (at, 1H, $J$ = 9.0 Hz, $H_{4'}$), 3.46 (dd, 1H, $J$ = 9.0 Hz, 2.9 Hz, $H_2$), 3.44 (dd, 1H, $J$ = 9.0 Hz, $J$ = 2.9 Hz, $H_{2'}$), 2.40-2.33 (m, 2H, $H_b$), 1.79 (m, 1H, OH), 1.70-1.19 (m, 26 H, $H_c$-$H_o$), 0.90 (t, 3H, $J$ = 6.7 Hz, $H_p$), 0.19-0.13 (6s, 54H, 6 $Si(CH_3)_3$); $^{13}C$ NMR ($CDCl_3$, 75 MHz); δ (ppm): 173.8 ($C_a$), 94.5, 94.4 (C1, C1'), 73.5, 73.3, 73.0, 72.8, 72.6, 71.9, 71.4 (C2, C2', C3, C3', C4, C4'), 70.8 (C5), 63.3 (C6), 61.6 (C6'), 34.2 (Cb), 32.0, 29.7, 29.7, 29.6, 29.5, 29.4, 29.3, 29.2, 24.8, 22.7 (Cc-Co), 14.1 (Cp), 1.1, 1.0, 0.9, 0.2, 0.1 (6 $Si(CH_3)_3$); ESI HRMS: calcd for $C_{46}H_{100}NaO_{12}Si_6$ $[M+Na]^+$: 1035.5723, found 1035.5718. Dowex 50WX8 ($H^+$ form) resin (457 mg) was added to a solution of the esterified compound (139 mg, 0.14 mmol) in MeOH (6 mL). The solution was stirred for 40 min at room temperature, then filtered, washed with MeOH and concentrated under reduced pressure. The residue was purified by flash silica gel chromatography ($CH_2Cl_2$/MeOH 90:10 to 80:20) to give the desired product **6** (78 mg, 96%) as a white solid. NMR data were in agreement with literature.[xi] $^1H$ NMR ($CDCl_3$/$CD_3OD$, 360 MHz); δ (ppm): 5.10 (d, 1H, $J$ = 3.7 Hz, $H_1$), 5.07 (d, 1H, $J$ = 3.7 Hz, $H_{1'}$), 4.36 (dd, 1H, $J$ = 2.1 Hz, $J$ = 11.9 Hz, $H_{6a}$), 4.20 (dd, 1H, $J$ = 5.1 Hz, $J$ = 11.9 Hz, $H_{6b}$), 4.02 (ddd, 1H, $J$ = 2.1 Hz, $J$ = 5.1 Hz, $J$ = 10.0 Hz, $H_5$), 3.85-3.74 (m, 4H, $H_3$, $H_{3'}$, $H_{6'a}$ and $H_{6'b}$), 3.67 ( m, 1H, $H_{5'}$), 3.49 (dd, 1H, $J$ = 3.6 Hz, $J$ = 9.7 Hz, $H_2$ or $H_{2'}$), 3.46 (dd, 1H, $J$ = 3.6 Hz, $J$ = 9.8 Hz, $H_2$ or $H_{2'}$), 3.33 (m, 2H, $H_4$ and $H_{4'}$), 2.34 (t, 2H, $J$ = 7.4 Hz, $H_b$), 1.61 (m, 2H, $H_c$), 1.38-1.23 (m, 24H, $H_d$-$H_o$), 0.9 (t, 3H, $J$ = 6.9 Hz, $H_p$); $^{13}C$ NMR ($CDCl_3$/$CD_3OD$, 90 MHz); δ (ppm): 175.3 (Ca), 94.5, 94.4 (C1, C1'), 73.9, 73.7, 73.1 (C3, C3', C5'), 72.5, 72.4 (C2, C2'), 71.3, 71.1 (C4, C4'), 70.7 (C5), 63.8 (C6), 62.2 (C6'), 34.7 (Cb), 32.5, 30.3, 30.2, 30.1, 30.0, 29.9, 29.8, 25.5, 23.3 (Cc-Co), 14.4 (Cp); ESI HRMS: calcd for $C_{28}H_{52}NaO_{12}[M+Na]^+$: 603.3351, found 603.3368.

#### 2- Supplementary data:

**Supplementary Table 1**

| | cMytC-TMM C13:0 |
|---|---|
| **X-ray source** | PROXIMA2 (2021/10/13) |
| **Wavelength (Å)** | 0.98013 |
| **Unit-cell (Å, °)** | 168.12, 168.12, 168.12, 90.0, 90.0, 90.0 |
| **Space group** | F23 |
| **Resolution limits (Å)** | 48.53 - 2.69 (2.85 – 2.69) |
| **Total reflections** | 107272(15776) |
| **Unique reflections** | 11045(1723) |
| **R-mean (%)** | 19.8(244.9) |
| **Completeness (%)** | 99.6(97.6) |
| **Mean I/σ (I)** | 12.21(1.09) |
| **CC (1/2)** | 99.7(34.9) |
| **Wilson B factor ($Å^2$)** | 68.4 |
| **Number of non-hydrogen atoms (protein/other)** | 2524/38 |
| **$R/R_{free}$ (%)** | 25.52/29.22 |
| **R.M.S.D. Bonds (Å)/angles (°)** | 0.007/0.93 |
| **Average temperature factors (protein/other)** | 86/96 |
| **Ramachandran plot (%) (Favored/Outliers)** | 93/1 |
| **PDB** | 8QHF |

**Supplementary Table 1: Data collection and refinement statistics**.
Statistics for the highest-resolution shell are shown in parentheses.

Supplementary Figure 1

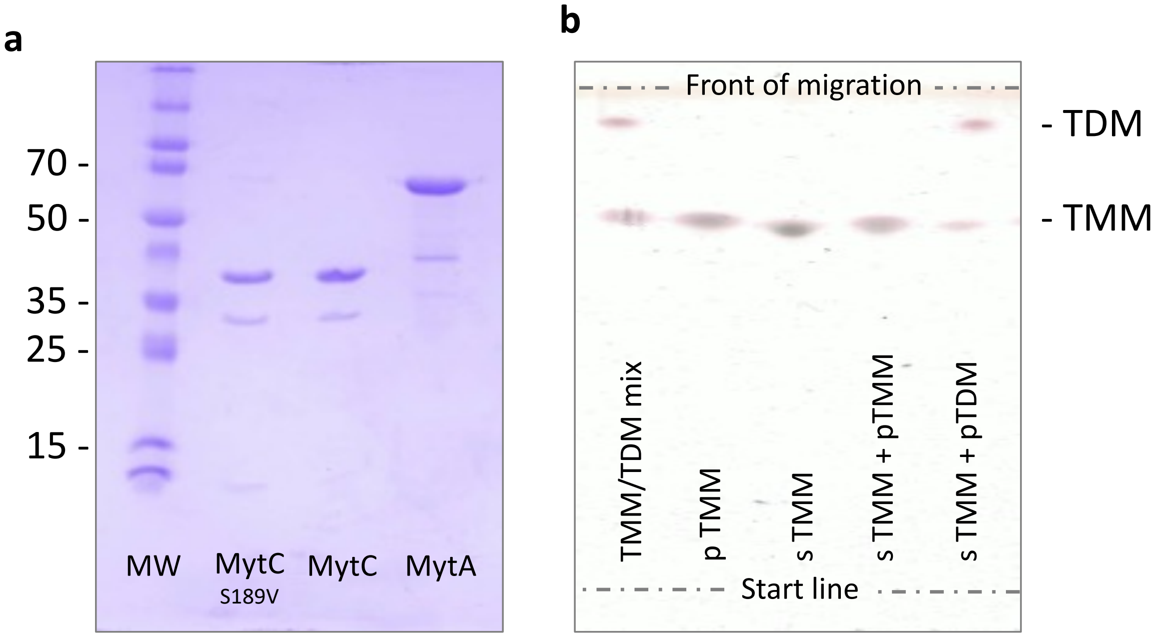


**Supplementary Figure1. Analysis of the enzymes and substrate donnors purified from bacteria.** (a) SDS/PAGE 12 % analysis of purified $MytC_{his}$ (WT), $MytC_{his}$ (S189V) and $MytA_{his}$, visualized by Coomassie blue coloration; (b) TLC of extractable lipids and synthetic TMM developed in $CHCl_3$-$CH_3OH$-$H_2O$ (34:15:2 vol/vol/vol) revealed by sulfuric acid staining.

**Supplementary Figure 2**

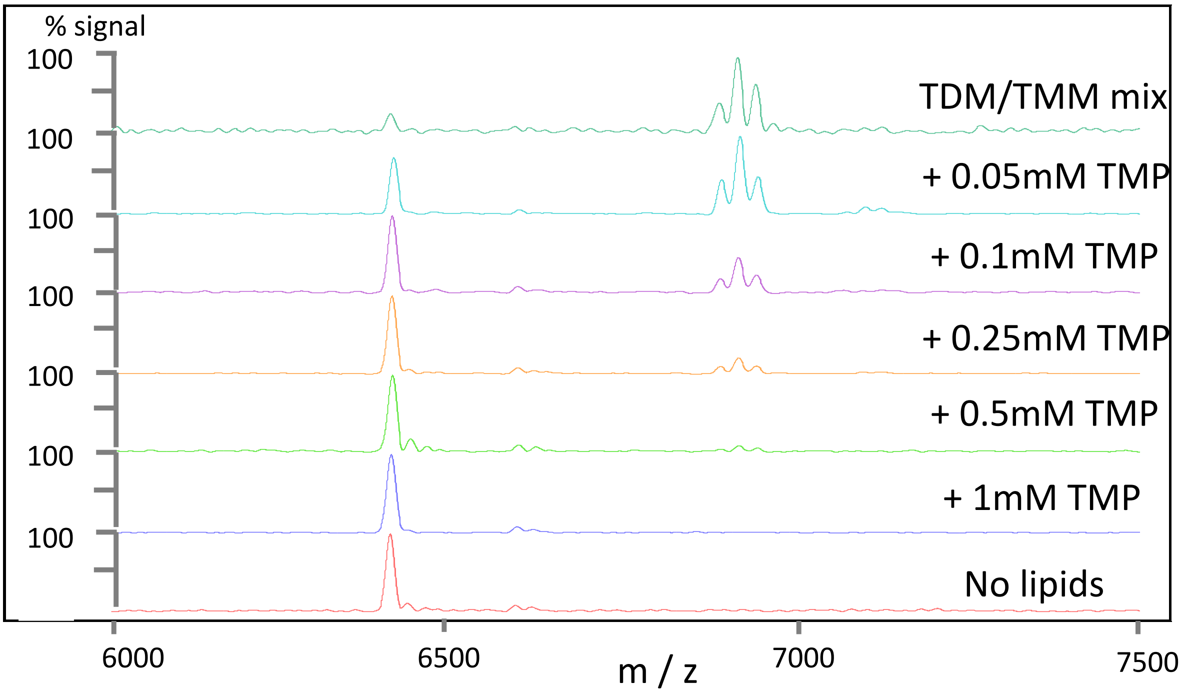


**Supplementary Figure 2. MytC inhibition by TMP.** Analysis of PorA mycoloylation in the presence of MytC and increasing TMP concentration (0.05 to 1 mM) by MALDI-TOF MS.

Supplementary Figure 3

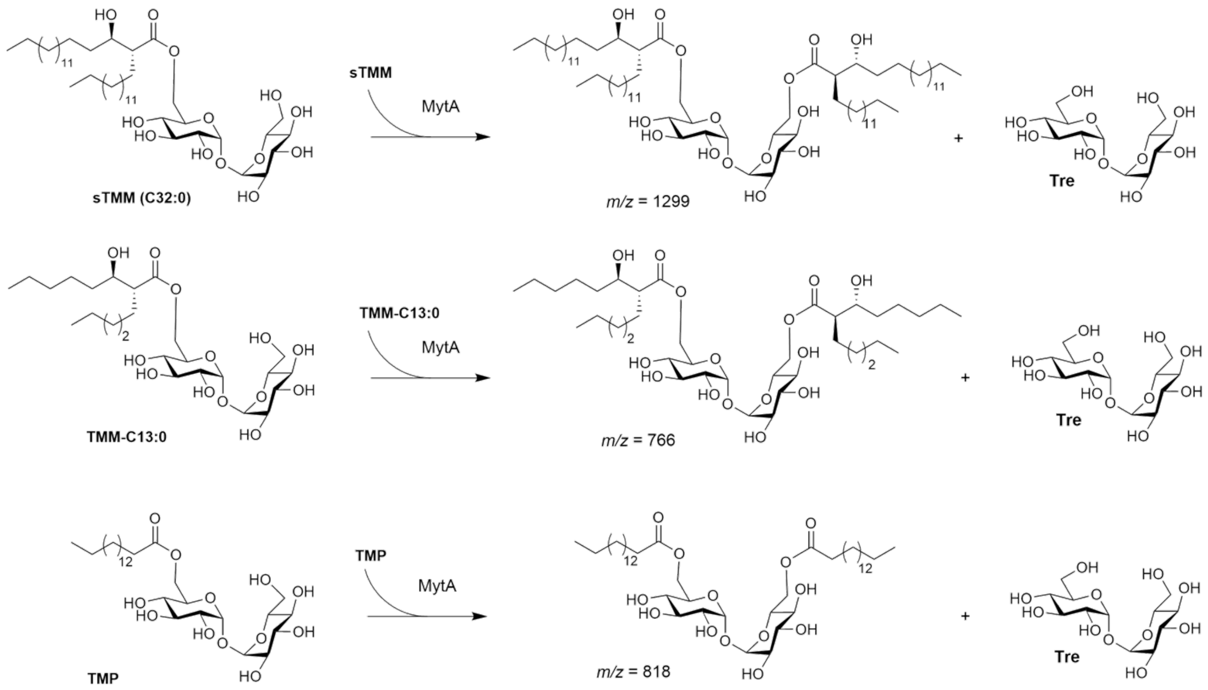


**Supplementary Figure 3. MytA processing of sTMM and analogs.** Structures and mass of the expected di-mycoloylated or di-acylated trehaloses products

Supplementary Figure 4

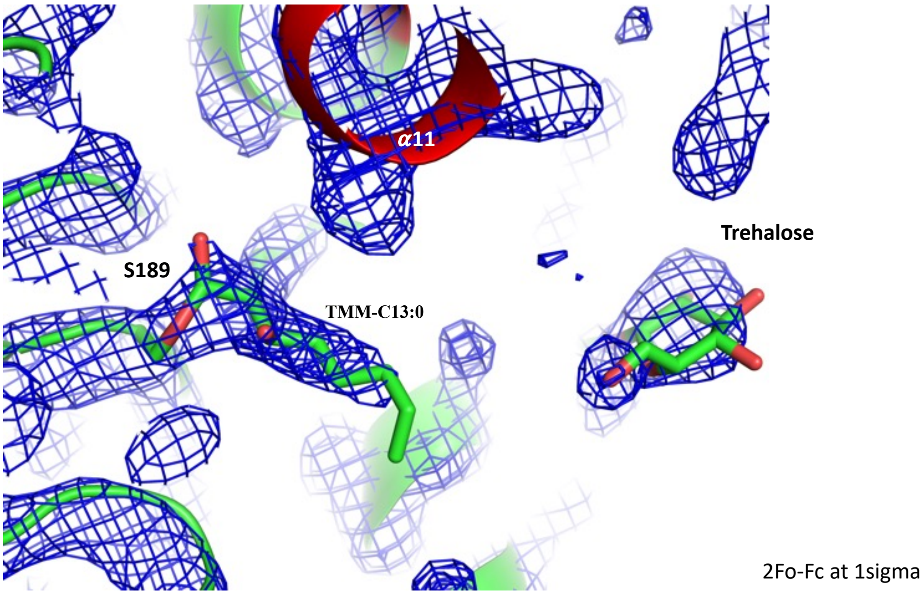


**Supplementary Figure 4. Crystal structure of the MytC-acyl enzyme**. 2Fo-Fc residual electron density around the covalently bound TMM-C13:0 contoured at 1 sigma.

Supplementary Figure 5

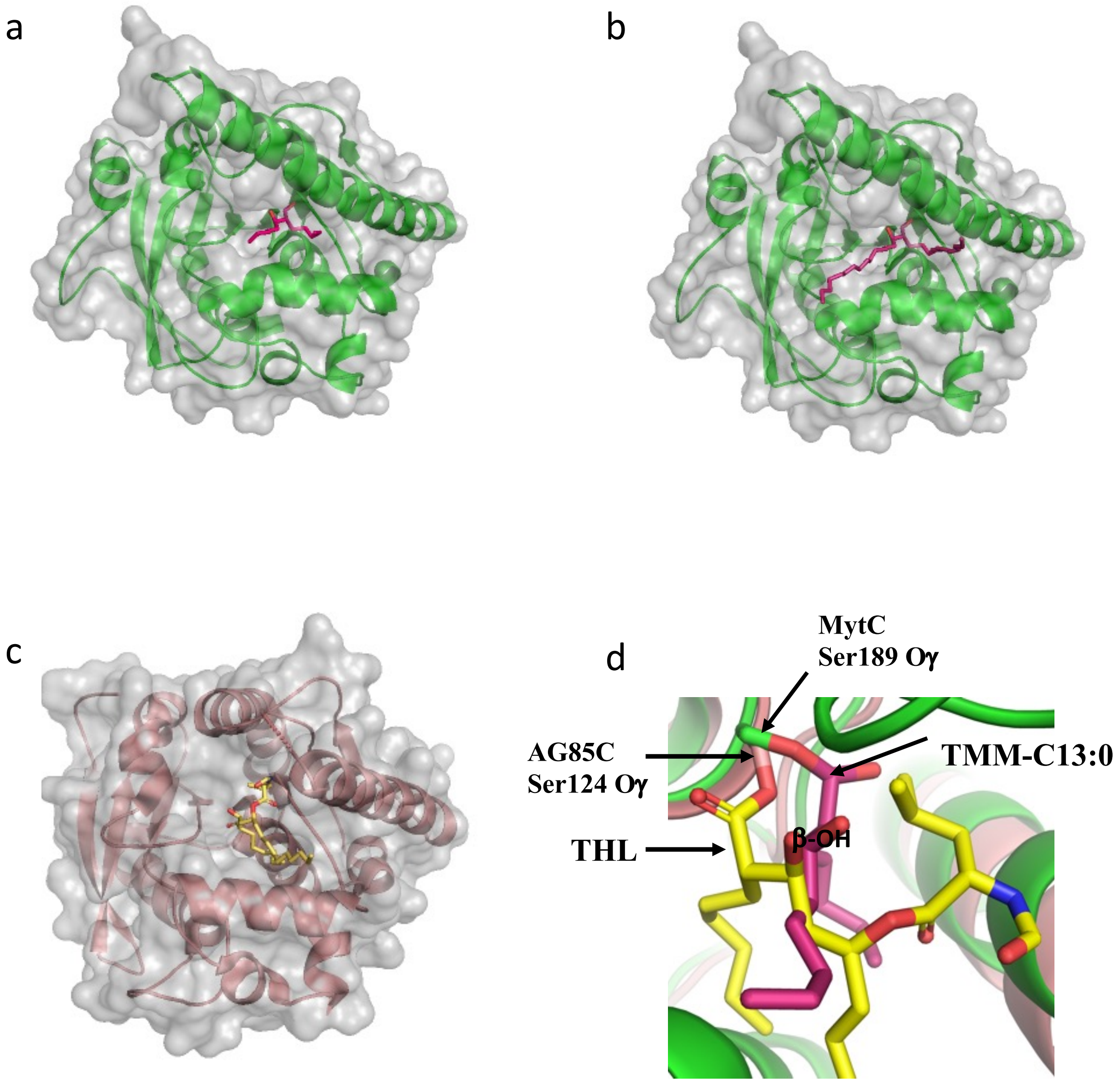


**Supplementary Figure 5. Alkyl pockets of mycoloyltransferases** (ribbon presentation imbedded in molecular surface). (a) MytC-acyl-enzyme : crystal structure of MytC covalently bound to TMM-C13:0 (red sticks) (b) Model of a MytC-acyl-enzyme bound to a natural mycoylate (red sticks). The model was obtained by replacing the short alkyl chains of TMM-C13:0 by long chains. (c) Ag85C-acyl-enzyme obtained with THL (sticks) (pdbcode 1VA5) Ag85C is shown in the same orientation as MytC in panels (a) and (b). (d) detailed view of the superposition of the acylated serines from the MytC and Ag85C acyl-enzymes. The β-OH position of both inhibitors (THL and TMM-C13:0) is also indicated.

#### Supplementary Figure 6

a b

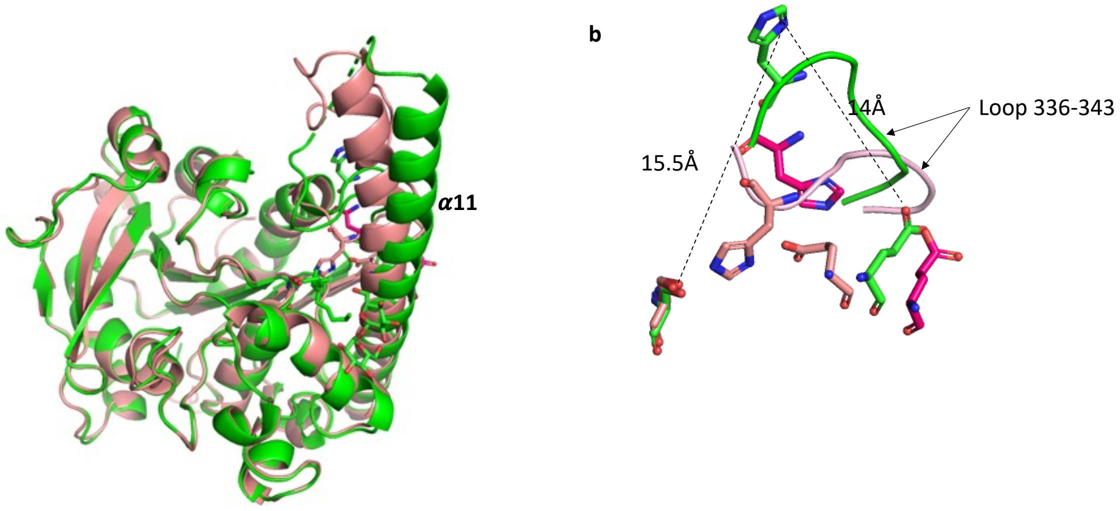


**Supplementary Figure 6. MytC structural changes between the apo and acyl-enzyme intermediate forms**. (a) Superposition of MytC model AlphaFold in wheat and MytC-acyl-enzyme in green; (b) MytC catalytic triad residues: MytC model AlphaFold in wheat, MytC apo enzyme in magenta, MytC-acyl-enzyme in green showing the histidine displacement.

**Supplementary Figure 7**

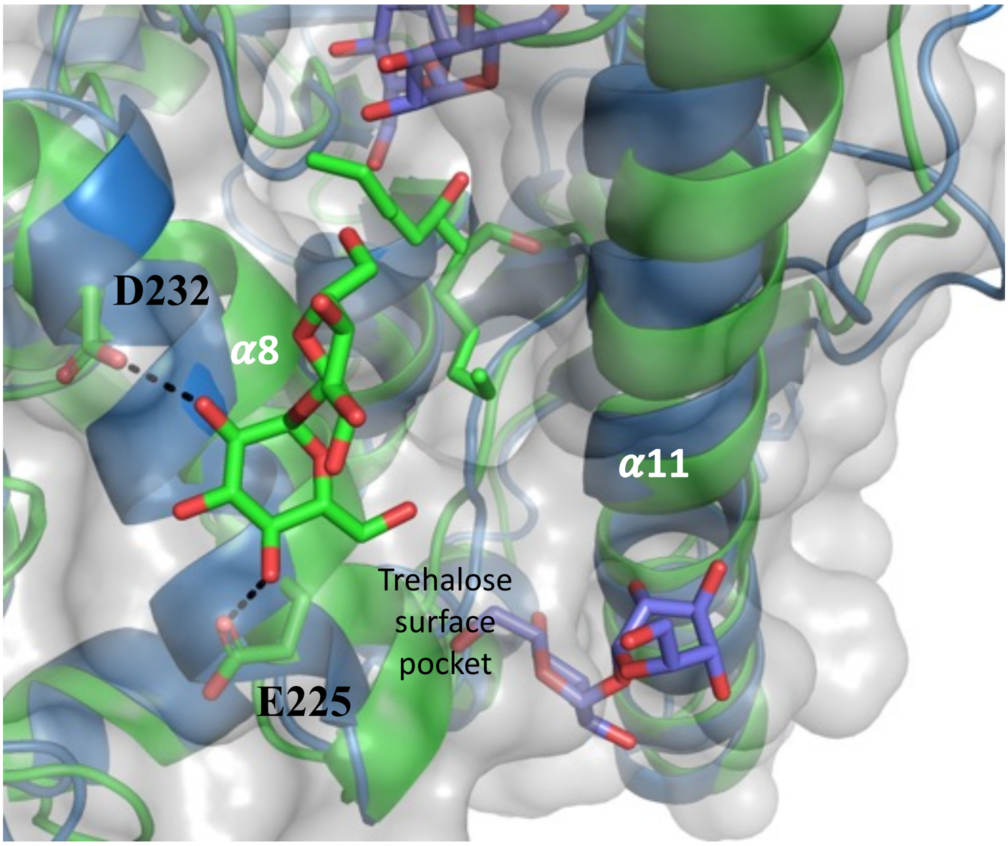


**Supplementary Figure 7**. **Trehalose binding site.** Surface pocket of the trehalose binding site. The MytC is in green and the Ag85 is in palecyan.

**Supplementary Figure 8**

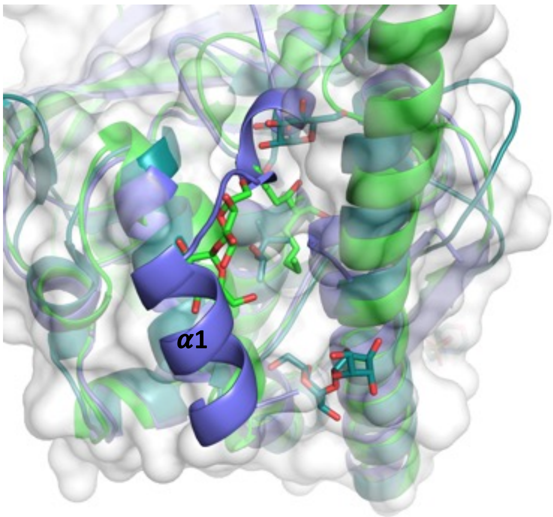


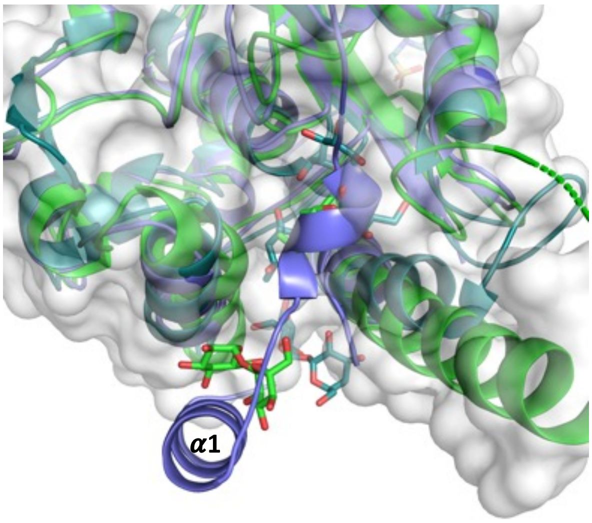


**Supplementary Figure 8. MytA/MytC and Ag85C superposition.** The α1 helix of MytA partially superposes with the PorA binding site of MytC and with a trehalose binding site of AG85C. The MytC is in green, the Ag85 is in palecyan and the MytA is in slate.

### Supplementary Figure 9

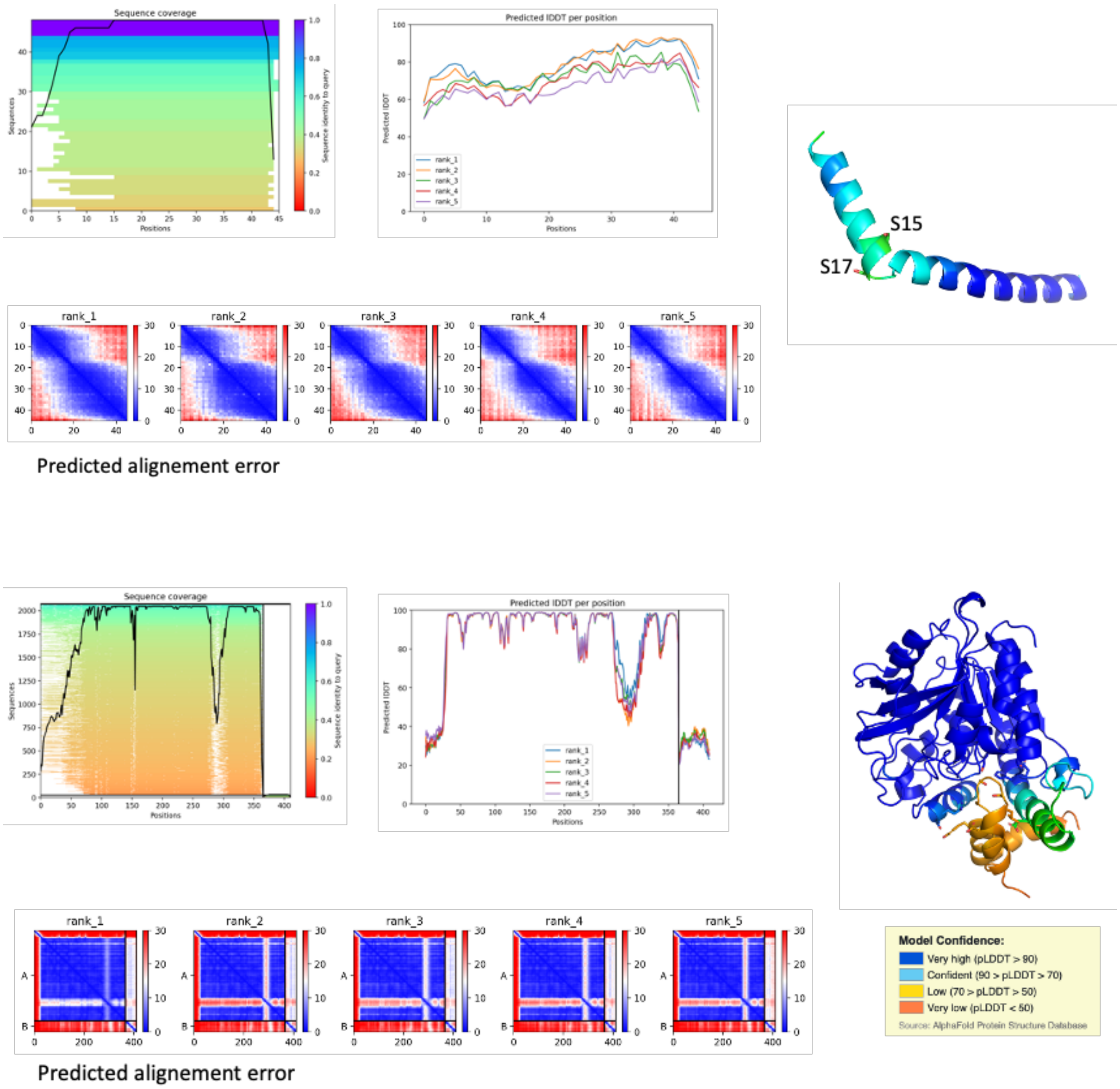


**Supplementary Figure 9. Statistic of AlphaFold model prediction for PorA (a) and MytC / PorA complex (b).** AlphaFold-v2; using version 2.3 of AlphaFold as implemented in our institute I2BC.

### Supplementary Figure 10

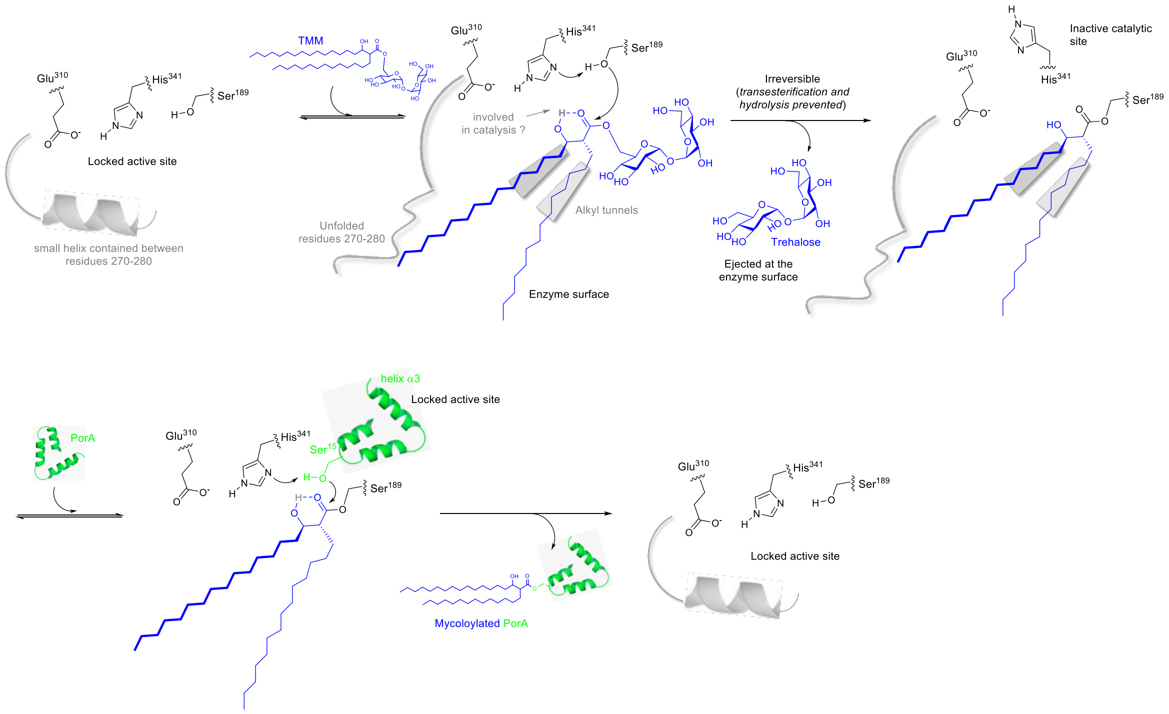


**Supplementary Figure 10. Proposed structure-based catalytic cycle of MytC**

[i] F. Migliardo, Y. Bourdreux, M. Buchotte, G. Doisneau, J.-M. Beau, N. Bayan, *Chem. Phys. Lipids*, **2019**, *223*, 104789.
[ii] P. L., Van der Peet, C. Gunawan, S. Torigoe, S. Yamasaki, S. J. Williams, *Chem. Commun*. **2015**, *51*, 5100-5103.
[iii] V. Ratovelomanana-Vidal, C. Girard, R. Touati, J. P. Tranchier, B. Ben Hassine, J. P. Genêt, *Adv. Synth. Catal*. **2003**, *345*, 261-274.
[iv] H. Yamamoto, M. Oda, M. Nakano, N. Watanabe, K. Yabiku, M. Shibutani, M. Inoue, H. Imagawa, M. Nagahama, S. Himeno, K. Setsu, J. Sakurai, M. Nishizawa, *J. Med. Chem.*, **2013**, *56*, 381-385.
[v] D. W. Brooks, L. D.-L. Lu, S. Masamune, *Angew. Chem. Int. Ed. Engl.*, **1979**, *18*, 72-74.
[vi] T. Seifert, M. Malo, T. Kokkola, E. J. L. Stéen, K. Meinander, E. A. A. Wallén, E. M. Jarho, K. Luthman, *Bioorganic & Medicinal Chemistry*, **2020**, *28*, 115231.
[vii] S. Muto, K. Mori, *Biosci. Biotech. Biochem.*, **2003**, *67*, 1559-1567.
[viii] J. P. Genêt, C. Pinel, V. Ratovelomanana-Vidal, S. Mallart, X. Pfister, M. C. C. De Andrade, J. A. Laffitte, *Tetrahedron: Asymmetry*, **1994**, *5*, 665-674.
[ix] J. Radivojevic, S. Skaro, L. Senerovic, B. Vasiljevic, M. Guzik, S. T. Kenny, V. Maslak, J. Nikodinovic-Runic, K. E. O'Connor, *Applied Microbiology and Biotechnology*, **2016**, *100*, 161-172.
[x] R. Toubiana, B. C. Das, J. Defaye, B. Mompon, M.-J. Toubiana, *Carbohydr. Res.*, **1975**, *44*, 308-312.
[xi] R. S. Kallerup, H. Franzyk, M. L. Schiøth, S. Justesen, B. Martin-Bertelsen, F. Rose, C. M. Madsen, D. Christensen, K. S. Korsholm, A. Yaghmur, C. Foged, *Mol. Pharmaceutics*, 2017, **14**, 2294-2306.
